# Computational evolutionary approach to generate antimicrobial peptides

**DOI:** 10.64898/2026.07.26.738612

**Authors:** Timothy Harrison, Tianmeng Zhang, Meet Parmar, Praveen Praveen, Aydin Menekse, Kevion K Darmawan, Alastair J Sloan, Wei Xiang, Andrew Hung, Wenyi Li

## Abstract

As a consequence of the overuse of conventional antibiotics, there is currently an unprecedented increase in antibiotic resistance in newer generations of pathogenic bacteria. This growing problem has led scientists to discover novel medications that could potentially reduce the usage of antibiotics, such as antimicrobial peptides (AMPs). Recent study has demonstrated that pardaxin could bind deeply into the surface of a lipid model membrane and inhibit the growth of pathogenic bacteria, including *Staphylococcus aureus* and *Escherichia coli*. In this work, the antibacterial efficacy of pardaxin was extended further by performing selective *in silico* substitution, driven by deep learning, Evolutionary-Scale Cambrian (ESMC) combined with conventional AMP design principles. Principal component analyses of the ESMC embeddings combined with conventional design models produced a set of single- and multiple-mutant analogues of pardaxin, which were predicted and validated with experimental data to exhibit selective antimicrobial properties against either *E. coli* or *S. aureus,* respectively. Atomistic molecular dynamics simulations further supported the notion that alpha-helical stability is a critical predictor of inner membrane activity, strongly correlated with their selective antimicrobial action tested in the lab. Overall, the findings highlighted a promising application of deep evolutionary machine learning techniques for screening a range of novel AMPs for selective antimicrobial agents.

## 1. Introduction

Antibiotic resistance has become a prevalent issue in global society that leads to the emergence of superbugs, including methicillin-resistant *Staphylococcus aureus* [1]. It is estimated that there could be 10 million deaths by 2050 [2]. Therefore, there is an urgent need for the development of new ways to curb this issue. Antimicrobial peptides (AMPs) are alternative therapeutics that have been shown to possess broad-spectrum antimicrobial activity with a lower tendency to develop antimicrobial resistance [3–6]. AMPs were originally thought only to be membrane disruptors, with their alpha-helical structure forming pores in the bacterial membrane causing cell lysis [7, 8]. Recently, it has become clear that they can also function as metabolic or ribosomal inhibitors, binding to key metabolic or ribosomal proteins and causing bacteriostasis [3, 8]. The traditional antibiotics can also be chemically cross-linked into these peptides, thereby improving their effectiveness against various pathogenic bacteria [9]. Thus, AMPs are considered as alternative therapeutics, diversifying infection treatments and reducing multidrug resistance.

Molecular modelling and simulations have been commonly used in the area of drug discovery and protein science as structural and activity-based techniques [10, 11]. They have been proven to be useful tools in obtaining atomic insights regarding the structural stability of proteins and their peptide segments under various physiological conditions [12–14]. For example, our recent study has applied molecular dynamics (MD) simulations to show the correlation of AMP Pardaxin’s antimicrobial activities with their structural ability for penetrating lipid membranes model of both Gram-positive and Gram-negative bacteria [15, 16]. On the other hand, Principal component analysis (PCA) has been used extensively in the analysis of molecular trajectories, feature extraction and statistical analysis [17], which can generate new protein descriptors in protein science. So far, there have been limited attempts made to predict minimum inhibitory concentration (MIC) through a PCA methodology using physiochemical descriptors, specifically for pardaxin.

Conversely, evolutionary guided algorithms and neural networks have been used extensively to predict the activity of AMPs [8, 18, 19]. Through an extended expression of physio-chemical descriptors, these computational approaches have enhanced AMP discovery by enabling the systematic exploration of vast sequence spaces that would be experimentally prohibitive to investigate through traditional methods. However, with these methods, it is difficult to elucidate the underlying characteristics of an active peptide. With classifier neural networks, although there is a proverbial ‘black box’ which produces predictions, the evaluating tangible design foundations from these predictions remains elusive. These methods obfuscate underlying structure-activity relationships and chemical pathways, making it difficult to extract meaningful design decisions. With clustering techniques, we can quickly evaluate a potential sequence’s efficacy, and those close interrogation of surrounding sequences, we can crystallise meaningful AMP design decisions for future AMP development.

Together with MD, PCA and neural networks, we aim to further elucidate the finding of Pardaxin interaction with lipid membranes by applying *in silico* techniques to iteratively induce selective substitutions onto Pardaxin for the purpose of fabricating novel Pardaxin-based peptides with enhanced antimicrobial efficacy and potential selectivity on different bacterial membranes.

## 2. Results

### 2.1 *In silico* deep learning-guided identification of prospective pardaxin mutants

To initially explore the extent to which PCA analysis of physicochemical descriptors can cluster AMP sequences, a set of 112 physicochemical descriptors was calculated, scaled and input to principal component analysis. MIC values reported from the database DBAASP for sequence variants with identity of greater than 0.8 for magainin-II, oncocin, pyrrhocoricin and peptide p were collected. Although the PCA analysis of magainin-II (DBAASPS_118) MIC values showed general activity trends (Figure 1), such as increasing PC1 and PC2 activity towards *E. coli*, it contains limited actionable clustering information for future AMP design. In addition, the PCA physicochemical descriptors provide non-actionable clustering formations (Figures 1A and 1D). Unlike traditional approaches that rely on predefined descriptors, the ESMC, an evolutionary neural network, represents a significant advancement in protein language models, trained on extensive protein sequence databases to capture evolutionary relationships and functional patterns embedded within amino acid sequences. Thus, ESMC was employed to provide a detailed embedding of a protein using 960 parameters across 30 layers. Indeed, Figures 1C and 1F showed a clear separation between non-active and active variants, particularly through layer 29, and thus, layer 29 was applied as the foundational layer of our search of active AMPs. The additional figures demonstrated the ESMC’s ability to cluster activity-structure relationships for the AMP variants of oncocin, pyrrhocoricin and peptide-p (Figure S3-S10).

**Figure 1.**
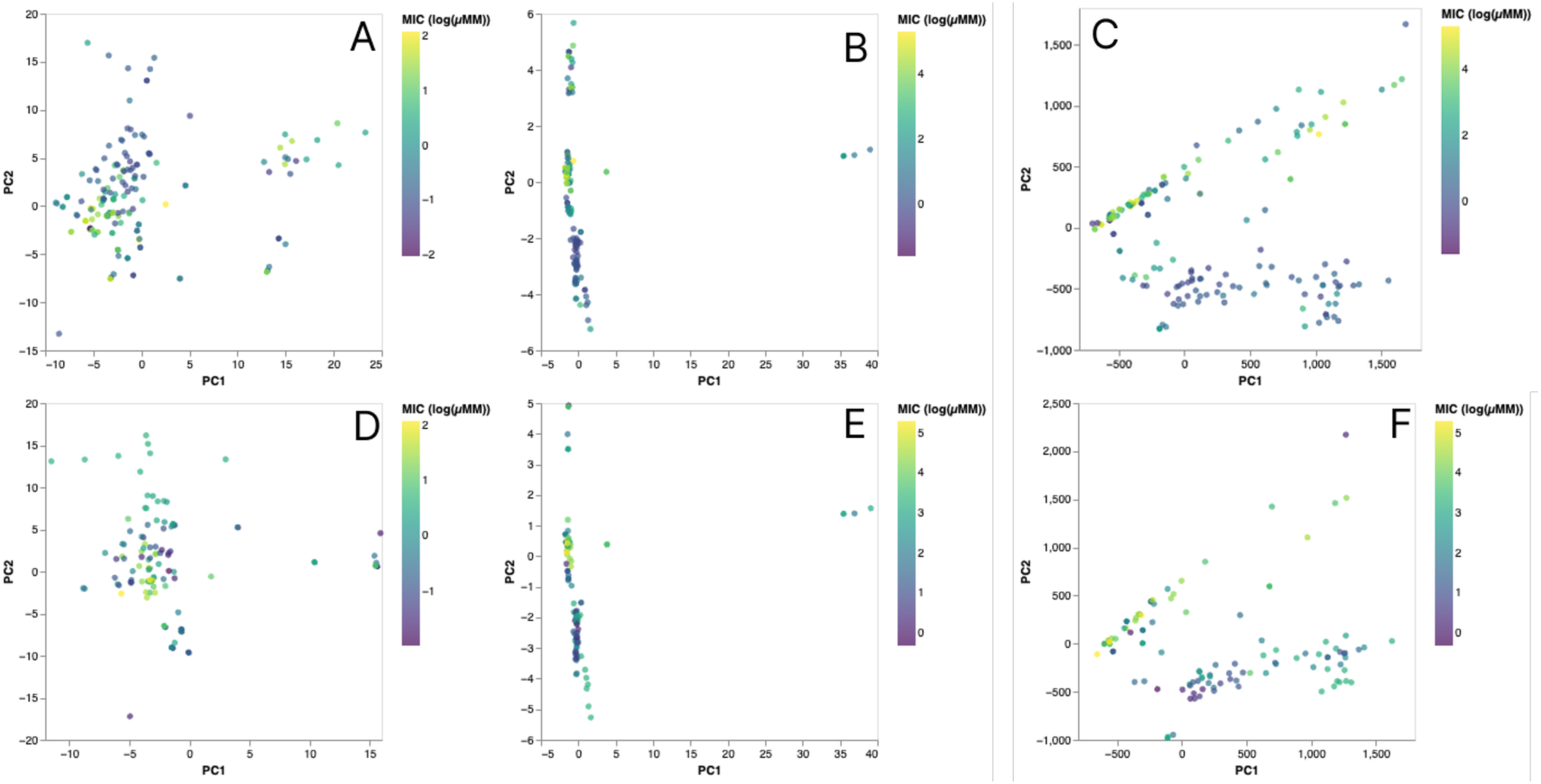
Scatter plots of PC1 and PC2 for magainin-2 variants with sequence identity > 0.8, and the base 10 logarithm of their MIC values. This was selected such that small MIC values were separated and large MIC values were collected. PCA of a set of 112 physicochemical descriptors and the MIC values against E. coli (A) and S. aureus (D). PCA of ESMC embeddings of layer 0 (B), and layer 29 (C) of the network, also coloured by Log (MIC) to E. coli. PCA of ESMC embeddings for layer 0 (E) and layer 29 (F), coloured by their activity towards S. aureus.

In the DBAASP database, there are only 12 reported pardaxin variants against *E. coli* and 5 pardaxin variants against *S. aureus* (accessed DBAASP in March 2026). To limit the initial search space, a comprehensive library of single-point substitutions was generated, systematically covering all 20 natural amino acids at each position of the 22-residue in pardaxin. By using ESMC and PC, Figure 2 showed layer 29 of the network embedding all the pardaxin substitutions, while overlaid are the known pardaxin variants from DBAASP and coloured with the logarithm of their MIC values. The MIC values are expanded in a range of 0 to 200, with the high activity are separated from the inactive sequences. To validate the effectiveness of ESMC PCA clustering on pardaxin, similar to that seen in magianin-II (Figure 1), points were selected across the distribution of the PCA space. Alpha-helicity, hydrophobicity and cationic nature were used in tandem to produce a set of variants for experimental evaluation (Figure 2).

**Figure 2.**
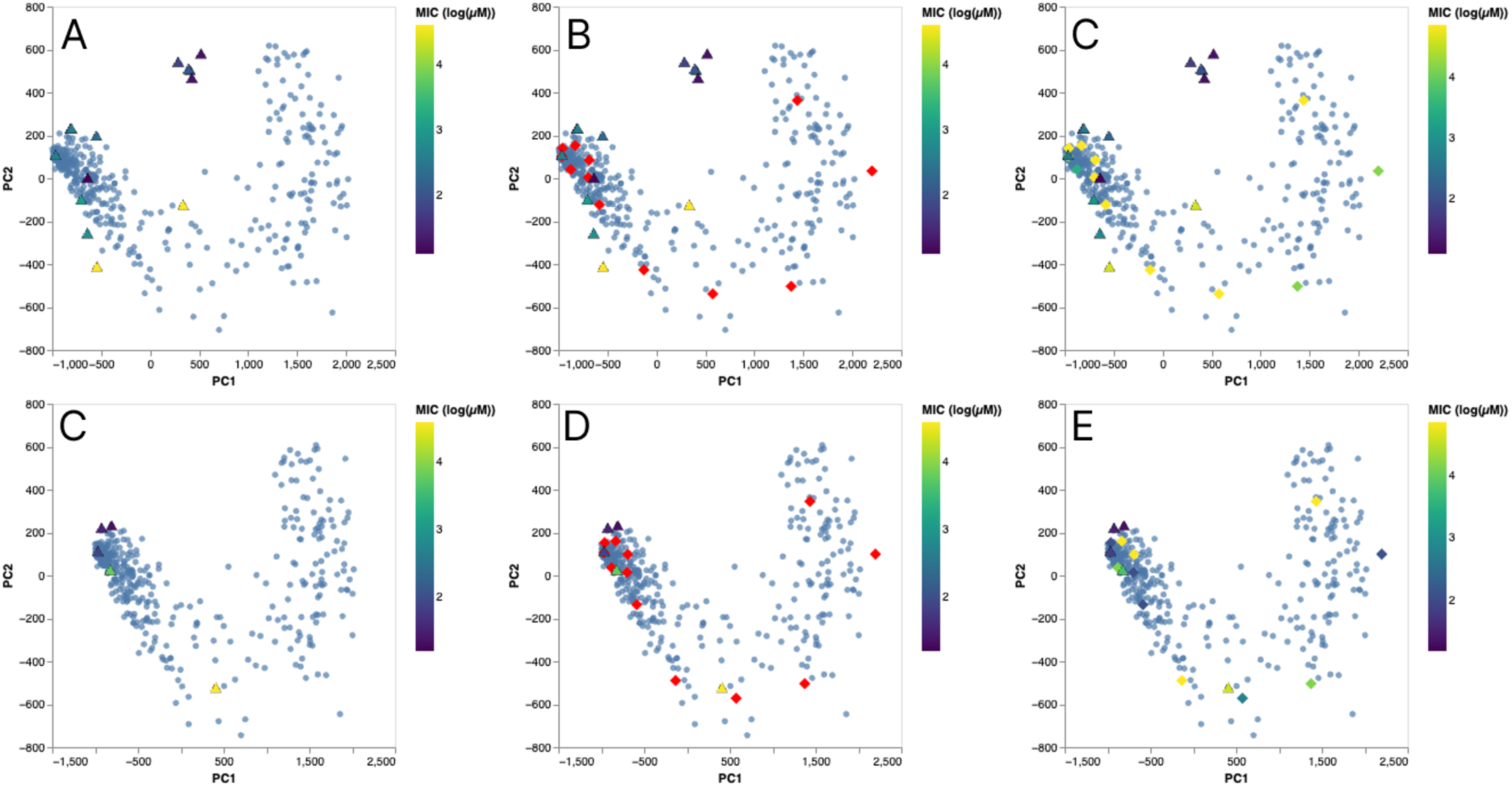
PCA1 and PC2 from the mean ESMC embeddings for layer 29 of pardaxin single-point substitution variants (blue). The distribution of active E. coli and S. aureus species is shown as coloured triangles, representative of the log of their MIC concentration, in (A) and (C), respectively. The distribution of the selected single-point substitutions, highlighting the distribution across the ‘arc’ of the PCA are shown in (B) and (D). The result of in vitro experimental testing is shown in (C) and (E) for E. coli and S. aureus respectively. Highlighting the ability of ESMC to infer structure-activity relationships through the clustering of active sequences at extremities of the distribution.

It has been shown that the addition of lysine to the hydrophobic face of AMPs can increase their potency. Particularly when substituting into hydrophilic positions increasing the amphicility, and thus, activity of the AMP [20–23]. To explore a multiple-point lysine substitution strategy, we substituted consecutive amino acids with Lysine, such that there are periodic faces of hydrophobicity and hydrophilicity, strengthening structure and producing hydrophobic faces. The resultant set of 14 Lysine substituted variants were mapped onto PCA space using their ESMC embeddings (Figure 3). Again, a combined selection profile was used: the distribution of the PCA was combined with the fact TPC122 and TPC123 are different only in the substitution of the Lysine at position 2 for TPC122 and position 3 for TPC 123, thus we also can examine the importance of the head “GFF” configuration of pardaxin (Figure 3 and their MIC values in Table 1).

**Figure 3.**
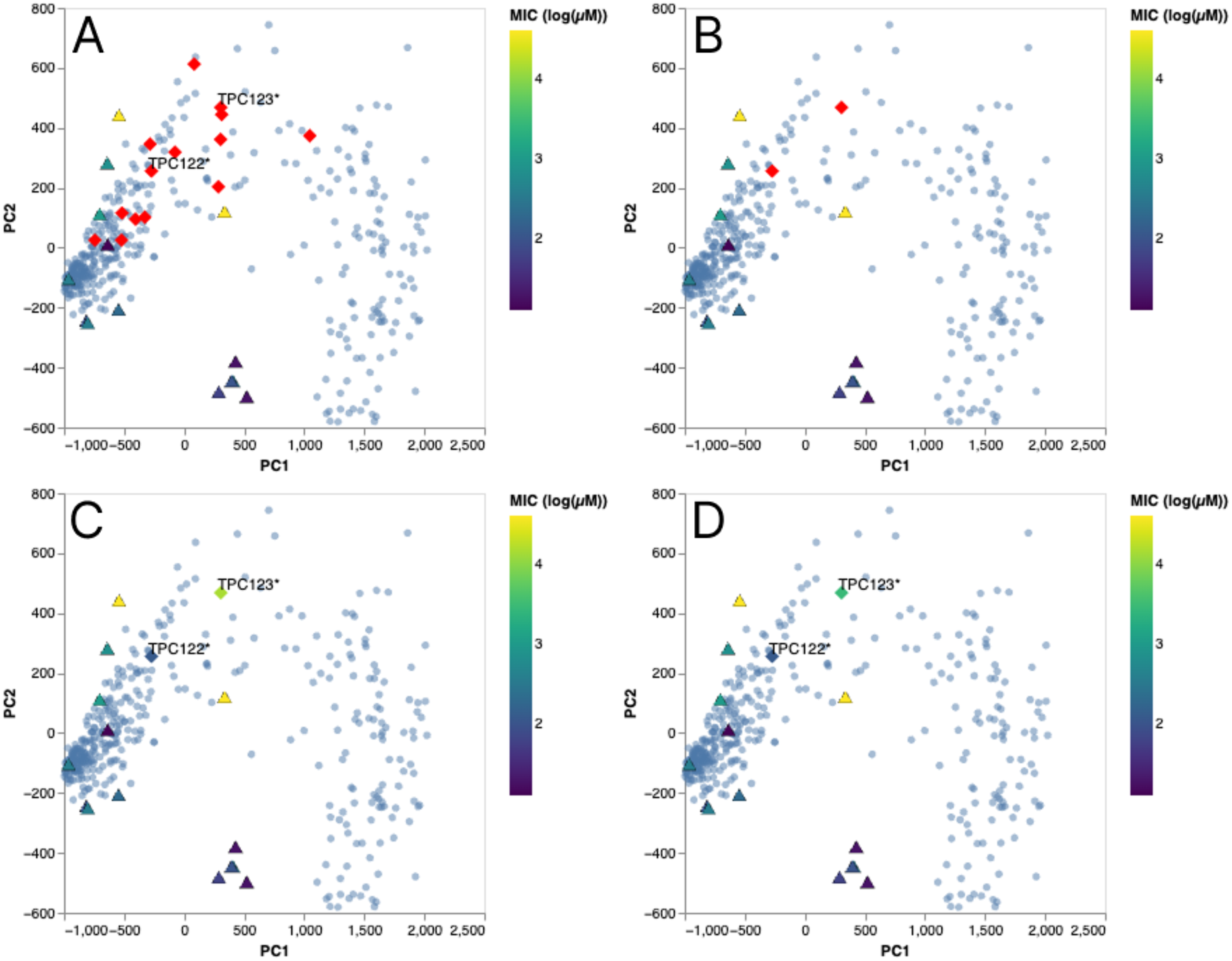
PCA1 and PC2 from the mean ESMC embeddings for layer 29 of pardaxin variants, the single-substituted sequences (blue), pardaxin reported activities from DBAASP against E. coli (colour gradient) and the set of lysine substituted sequences (red diamond) (A) and (B). Log of the experimental values of TPC123 and TPC122 against E. coli species, E. coli 25922 (C) and E. coli 35218 (D).

**Table 1.**
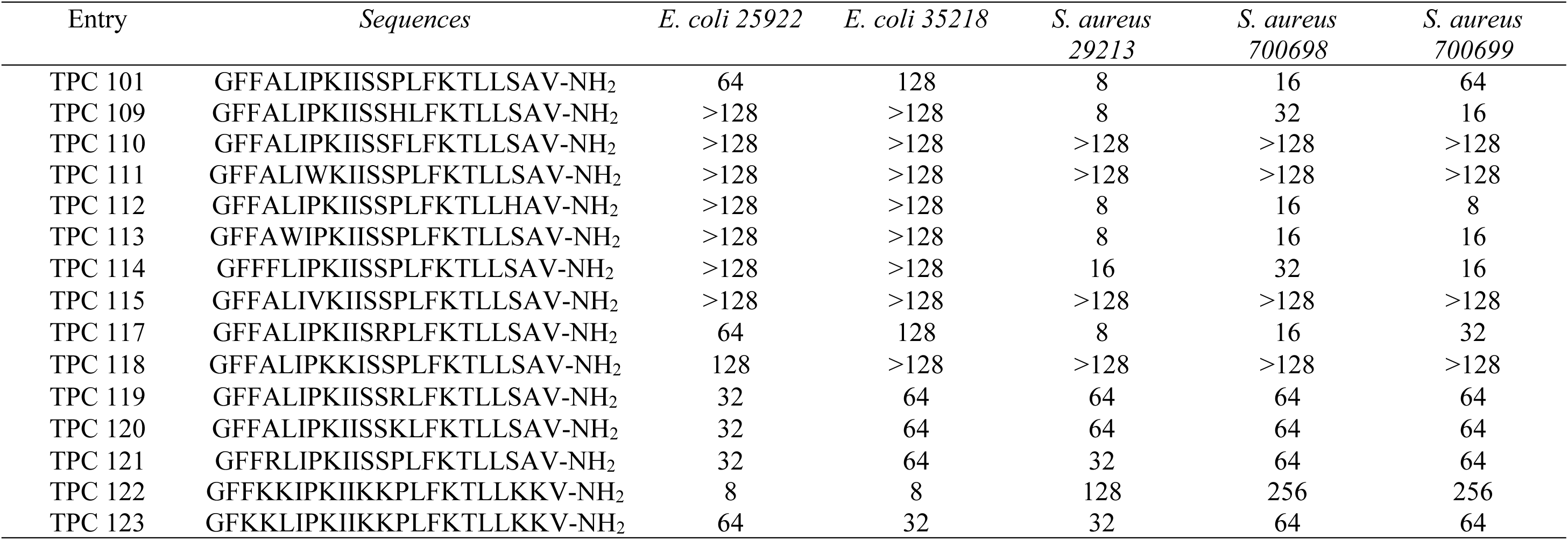
Antibacterial activity (MIC (µg/mL) of peptides against Gram-positive and Gram-negative bacteria

### 2.3 Antimicrobial activity of selected variants

The selected peptides (Table 1) were synthesised via standard Fmoc/tBu solid phase peptide synthesis as we previously described [15, 16, 24]. Their antibacterial activities were evaluated against representative Gram-negative (*E. coli 25922* and *35218*) and Gram-positive strains, including methicillin-sensitive and *methicillin-resistant Staphylococcus aureus* (MRSA). TPC101 (Pardaxin) and TPC113 preferentially inhibited Gram-positive bacteria, displaying strong activity against both methicillin-sensitive and MRSA strains, while exerting comparatively limited effects on Gram-negative *E. coli*. In contrast, TPC122 demonstrated pronounced activity against Gram-negative bacteria (8µg/mL) but only marginal efficacy against Gram-positive strains. TPC118 showed little to no antibacterial activity under the conditions tested. Together, these results highlight clear differences in antibacterial spectrum within the Pardaxin series, suggesting that subtle sequence variations confer distinct bacterial targeting properties, which correlates with the separation in Figures 1 and 2.

### 2.6 ESMC Clustering evaluation

Both TPC122 and TPC123 from Table 1 were shown to be selectively active against *E coli.* The position substitution motif at position 3 (TPC123) reported less activity against *E. coli* than the position substitution at 4 (TPC122). This provides more evidence that for AMP activity there is a strong correlation - not only between positive charges and activity - but specifically the periodic alternation of hydrophobic and positive sidechains. This is probably due to higher structural stability within the hydrophobic envelope of the membrane, as well as better adsorption onto the membrane, due to the positive charges.

Interestingly, the resultant grouping of the selected variants and their activities against *E. coli* species and variants against *S. aureus* species (Figure 2&3) strongly suggested the active variants to both *E. coli* and *S. aureus* species exhibiting areas of low PC1, and high PC2. Combined with the results from the DSAAB search of sequence similar peptides (Figure S9&S10), this demonstrates that ESMC has some capacity within its protein embedding system to capture the non-linear activation schemas of AMPs. This enables future development of AMP sequences through reinforcement learning practices.

### 2.5 Membrane-disruptive mechanisms of selected mutants

To further unveil the membrane activity of the selected Pardaxin variants, especially the active ones, TPC118 & TPC122 against *E. coli*, TPC113 & TPC118 against *S. aureus*, we further performed the membrane permeability for the *E. coli* outer/inner membrane and *S. aureus* cytoplasmic membrane, respectively.

#### a) Outer membrane permeabilisation in *E. coli*

Outer membrane integrity of *E. coli* was examined using the N-phenyl-1-naphthylamine (NPN) uptake assay (Figure 4). Exposure to TPC122 resulted in a rapid elevation of fluorescence signal, consistent with increased outer membrane permeability, with similar responses observed at other tested concentrations (Figure S11). TPC101 and TPC118 showed only minimal effects under the same conditions. Overall, TPC122 exhibited substantially greater outer membrane-disruptive activity than the other peptides, which showed limited effects across the tested range.

**Figure 4.**
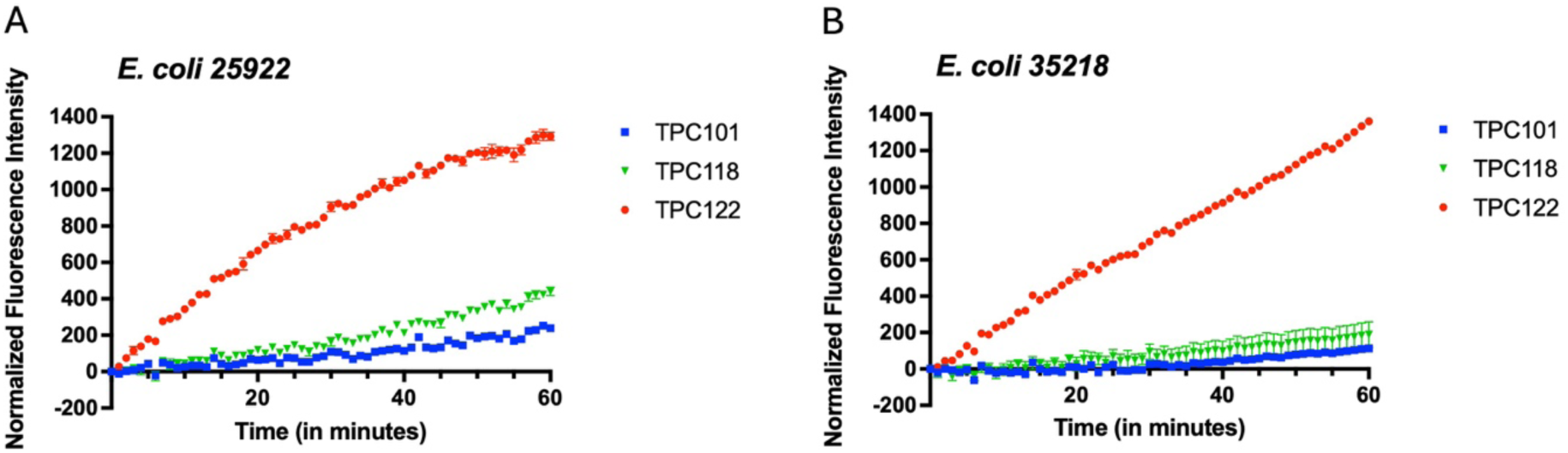
Peptide-induced outer membrane permeabilisation. NPN fluorescence kinetics were monitored to assess outer membrane permeabilisation following exposure to TPC101, TPC118, or TPC122 at concentration of 8 µg/mL. (A) E. coli 25922 and (B) E. coli 35218 were incubated with the indicated peptides, and fluorescence intensity was recorded over time. Enhanced NPN fluorescence indicates increased disruption of the bacterial outer membrane. Data are presented as mean ± SEM from three independent experiments.

#### b) Inner membrane permeabilisation in *E. coli*

Inner membrane integrity was assessed using the o-nitrophenyl-β-D-galactopyranoside (ONPG) hydrolysis assay, which reflects membrane disruption by permitting substrate access to intracellular β-galactosidase. At 8 µg/mL, TPC122 triggered a pronounced increase in ONPG hydrolysis in *E. coli*, consistent with effective inner membrane perturbation. By contrast, TPC101 and TPC118 elicited minimal changes under the same conditions (Figure 5, Figure S12). These results indicate that TPC122 substantially compromises the inner membrane of *E. coli*, whereas the other peptides have limited impact.

#### c) Cytoplasmic membrane disruption in *S. aureus*

The membrane-disruptive activities of TPC101, TPC113 and TPC118 against *S. aureus* were assessed by monitoring fluorescence over time (Figure 6, Figure S13). TPC101 induced a rapid increase in fluorescence, particularly within the first 10 min, followed by a plateau phase, indicating effective membrane permeabilisation. This effect became more pronounced with increasing peptide concentration, demonstrating a clear concentration-dependent response. TPC113 also displayed membrane-disruptive activity that was evident mainly at higher concentrations. In contrast, TPC118 produced minimal changes in fluorescence intensity across all tested concentrations.

**Figure 5.**
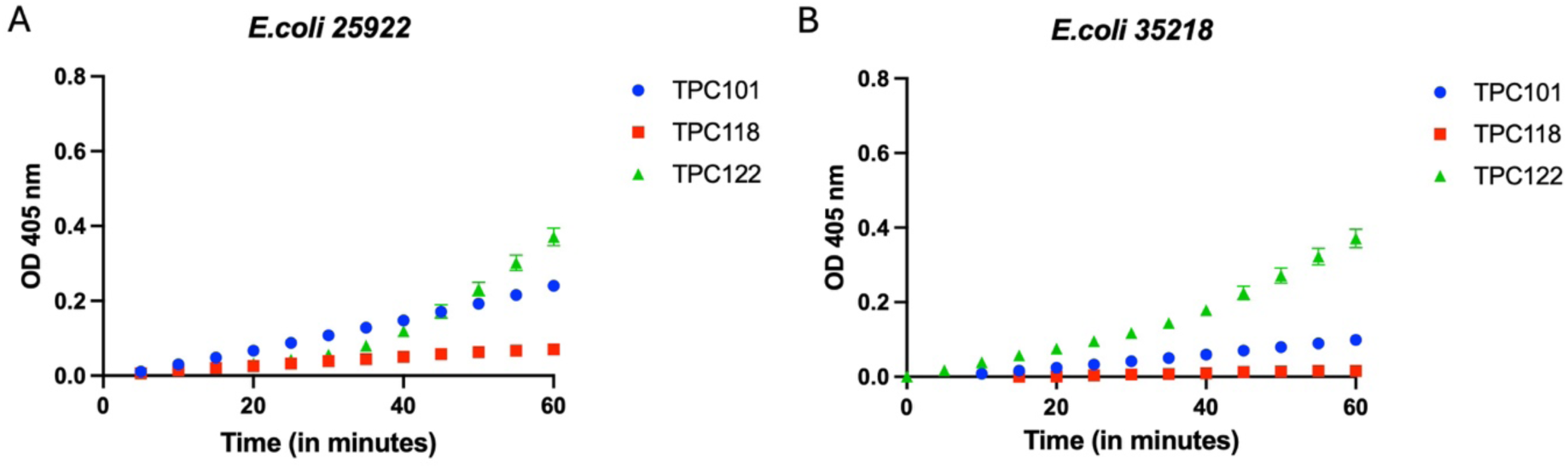
Peptide-induced inner membrane permeabilisation. ONPG hydrolysis was used to assess cytoplasmic membrane integrity following treatment with TPC101, TPC118, or TPC122 at a fixed concentration (8 µg/mL). (A) E. coli 25922 and (B) E. coli 35218 were incubated with the peptides, and o-nitrophenol formation was monitored spectrophotometrically over time. Increased absorbance indicates enhanced membrane permeabilisation, allowing ONPG access to intracellular β-galactosidase. Data represent mean ± SEM from three independent experiments.

**Figure 6.**
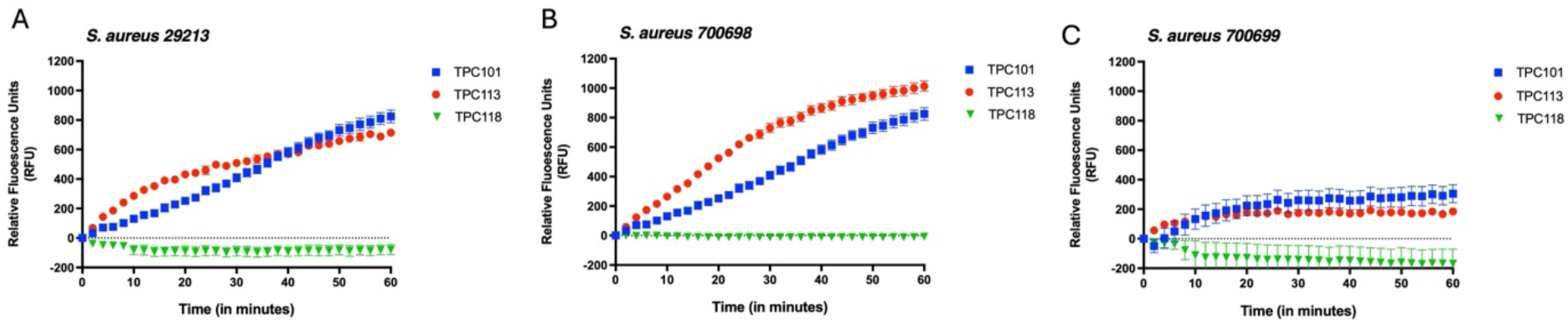
Peptide-induced cytoplasmic membrane permeabilisation of S. aureus. Membrane integrity was assessed by monitoring fluorescence over time following treatment with TPC101, TPC113, or TPC118 at concentration of 8 µg/mL. (A) S. aureus 29213, (B) S. aureus 700698, and (C) S. aureus 700699 were incubated with the peptides, and fluorescence intensity was recorded over time. Increased fluorescence reflects enhanced cytoplasmic membrane permeabilisation. Data represent mean ± SEM from three independent experiments.

Collectively, these results reveal distinct membrane-disruptive profiles among the peptides, with TPC101 exhibiting the strongest and most concentration-dependent activity, TPC113 showing moderate effects at elevated concentrations, and TPC118 remaining largely inactive.

### 2.4 Ultrastructural alterations in bacteria following peptide treatment

Their membrane activity was further validated with scanning electron microscopy (SEM) via marked morphological changes in both bacterial species following peptide treatment (Figure 7). *E. coli* exhibited smooth, intact rod-shaped cells with clearly defined boundaries in the control group. Upon TPC122 treatment, cells showed initial rupture and mild shape irregularities at lower concentrations, progressing to extensive cracking, elongation, and fragmentation at higher concentrations, indicating substantial membrane disruption (Figure 7A-4D). *S. aureus* maintained intact, spherical morphology in the control group; exposure to TPC113 caused surface wrinkling and rupture, with widespread breakage, loosening of intercellular connections, and accumulation of cellular debris at higher concentrations, culminating in near-complete lysis (Figure 7E-4H). These SEM observations highlight the disruptive effects of TPC122 and TPC113 on *E. coli* and *S. aureus*, respectively, reflecting their distinct impacts on bacterial membrane.

**Figure 7.**
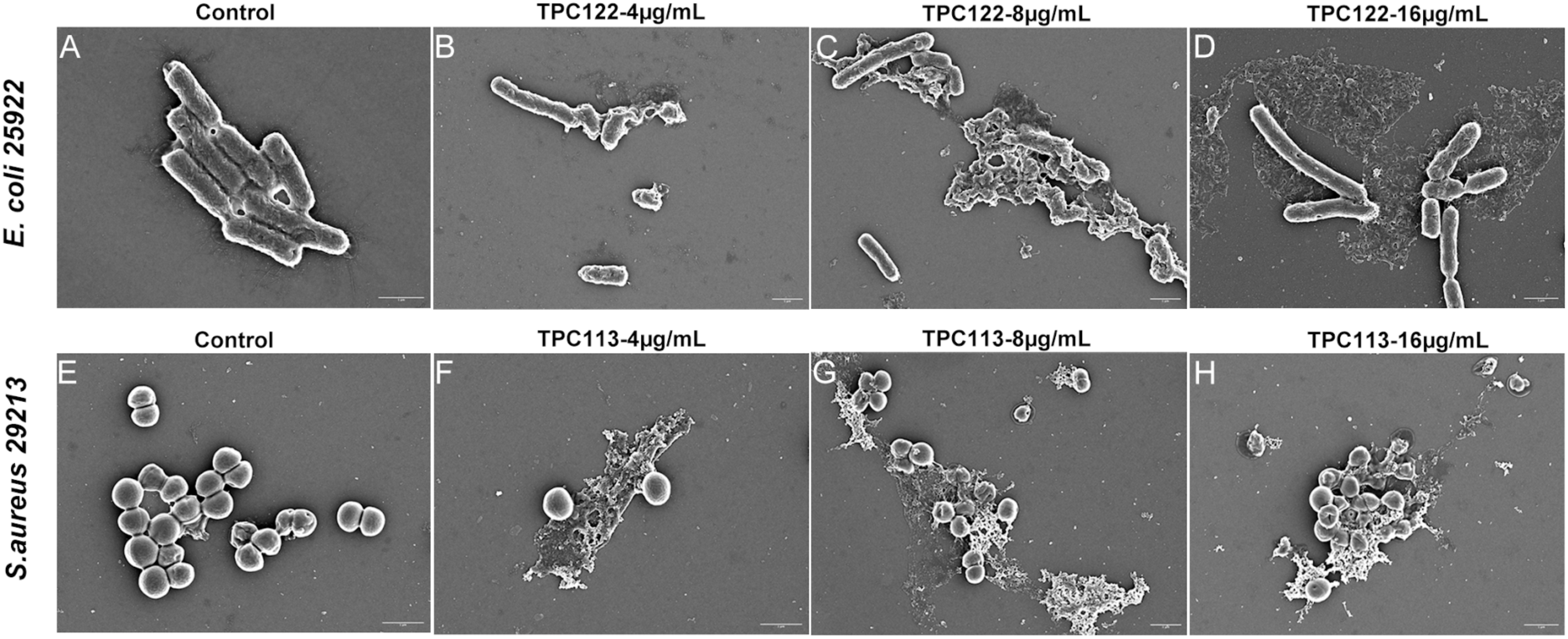
Scanning electron microscopy (SEM) analysis of bacterial morphology following peptide treatment. (A–D) E. coli 25922 and (E–H) S. aureus 29213 were either untreated (control, A and E) or exposed to peptides at 4, 8, or 16 µg/mL (B–D, F–H). Scale bar=1 µm.

### 2.2 Molecular dynamics evaluation of structural stability

To understand the dynamic process of Pardaxin variants’ selective interaction with Gram-negative and Gram-positive membranes, MD simulations were performed to elucidate the biophysical basis underlying the differential bioactivity of TPC113 and TPC122 toward two model membranes representing distinct bacterial inner membrane compositions, Gram-positive (9POPG:1POCL) and Gram-negative (7POPE:3POPG).

As shown in Figure 8A (blue line), TPC113 exhibited a slow and progressive increase in lipid contacts with the Gram-negative (7POPE:3POPG) membrane over the course of the 450 ns simulation, consistent with gradual surface association and stabilisation at the bilayer interface. The steady increase in contacts suggests electrostatic attraction to anionic POPG lipids followed by hydrophobic insertion within the membrane core. Such interaction dynamics revealed membrane-specific adsorption and secondary structure that correlate with experimentally observed activity differences. In contrast, for the Gram-positive (9POPG:1POCL) membrane, TPC113 at first maintained minimal contacts for approximately the first 300 ns of simulation, indicating limited early-stage affinity. Importantly, after ∼300 ns, the peptide underwent an abrupt transition characterised by rapid membrane engagement and insertion, resulting in a substantially higher number of lipid contacts than observed for Gram-negative membrane (Figure 8A, red line). This late insertion suggests the presence of a kinetic barrier to insertion in Gram-positive membrane that, once overcome, permits deeper bilayer integration (Figure 8C). Per-residue decomposition of each TPC113 residue in contact with all lipids of the NEG (Figure 8B, blue) and POS (orange) indicates markedly higher lipid contacts at the Gram-positive model membrane across the entire length of the peptide, marked with asterisks in the figure, indicating a broad, favourable interaction profile for Gram-positive relative to Gram-negative membranes. These simulation results are consistent with the low MIC for TPC113 at *S. aureus* relative to *E. coli* (Table 1).

**Figure 8.**
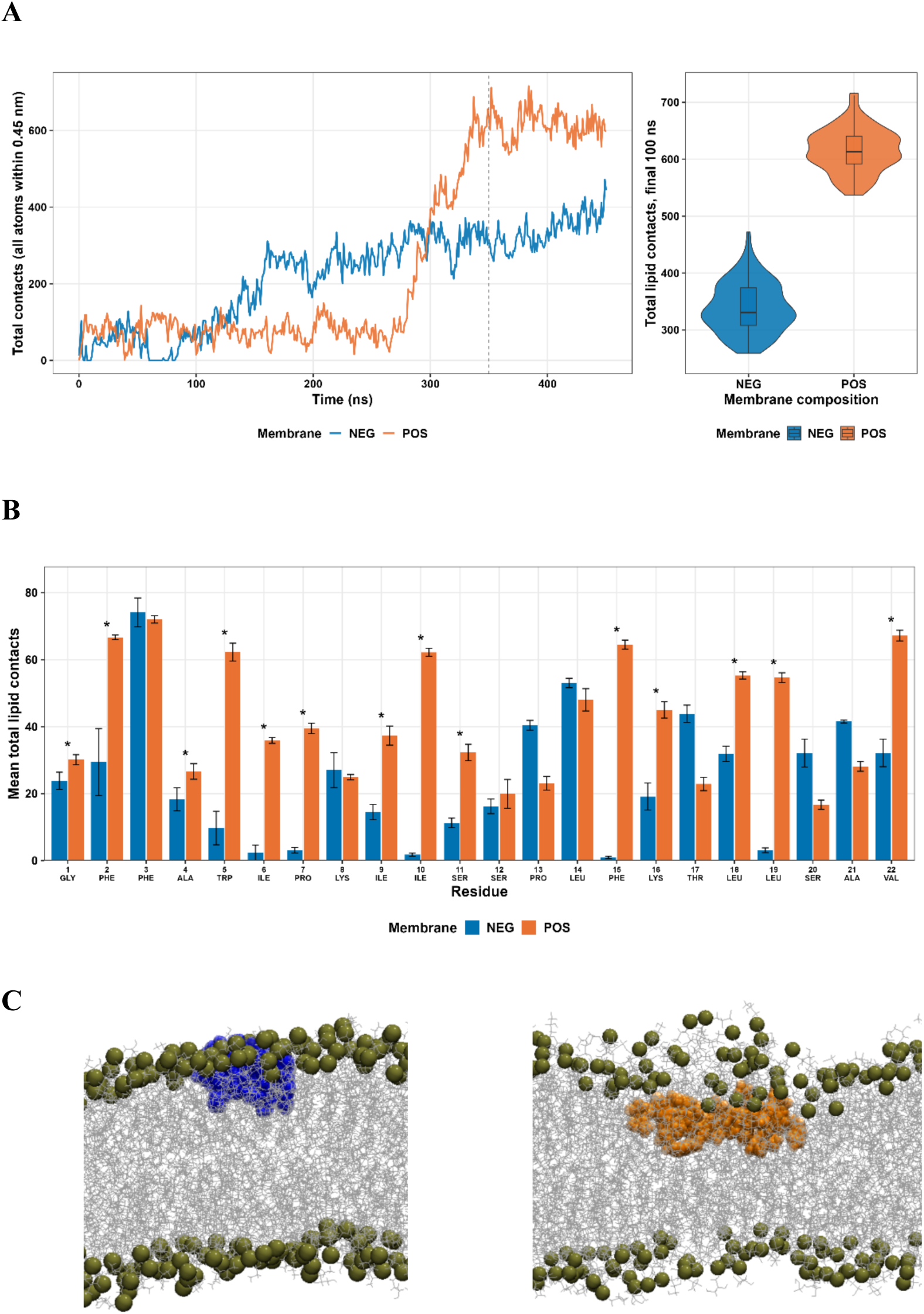
MD simulations of TPC113 at model Gram negative (NEG) and positive (POS) membranes. (A) Time series of interatomic contacts within 0.45 nm between TPC113 peptide and model NEG (blue) and POS (orange) membranes. (B) Per-residue bar charts of peptide-membrane contacts, with averages taken over the final 100 ns. Error bars are five-block-averaged standard errors over the relevant trajectory segment. (C) Graphical snapshots of TPC113 at NEG (blue) and POS (orange) membranes, with bronze spheres representing lipid phosphates, indicating the general extent of membrane insertion.

The TPC113 secondary structure analysis of α-helicity (ellipticity at 222 nm in the model system) was marginally greater in the presence of a Gram-positive membrane compared to a Gram-negative membrane (Figure 9A). A similar trend was shown by the simulation-derived percentage helicity profile, with more sustained helical content in the Gram-positive membrane, particularly at the N-terminal residues Gly1 to Lys8 (Figure 9B). Helicity is widely recognised as a key structural determinant of membrane-active peptide function, as α-helical conformations promote amphipathic structure and facilitate insertion into lipid bilayers. These findings are consistent with experimental results demonstrating greater biological activity of TPC113 against *S. aureus*.

**Figure 9.**
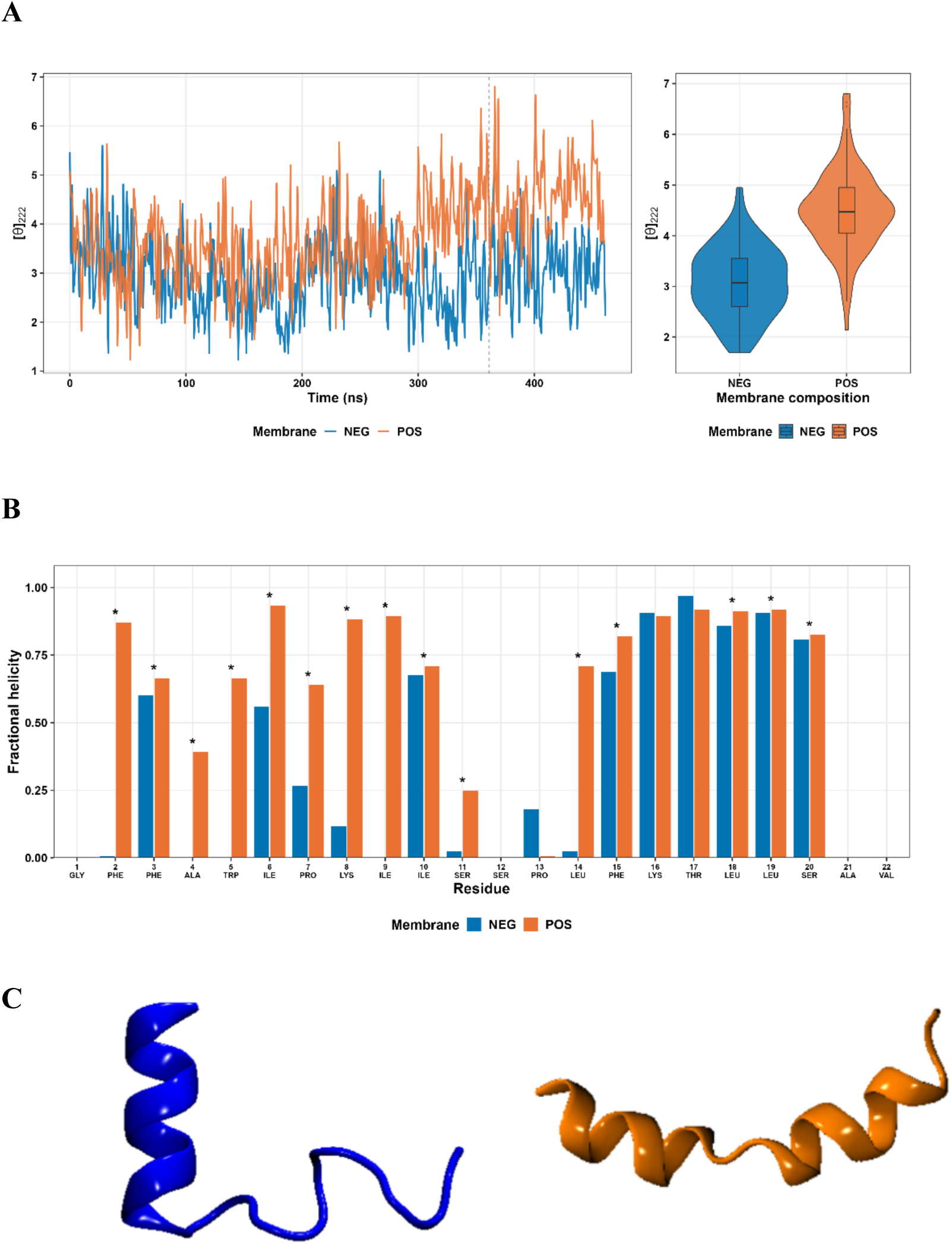
MD simulations of TPC113 at model Gram negative (NEG) and positive (POS) membranes. (A) Time series of ellipticity at 222 nm for TPC113, indicating helicity. (B) Per-residue bar charts of fractional helicity, averaged over the final 100 ns. (C) Graphical snapshots of TPC113 at NEG (blue) and POS (orange) membranes, indicating the general extent of helical content for the peptide at each membrane.

For TPC122, the relative trends in terms of membrane engagement and helical stability at Gram-positive and Gram-negative membranes are weaker, but still consistent with experiments. In contrast to TPC113, the TPC122 peptide displayed gradual increase in contacts, plateauing after around 200-250 ns (Figure 10A). While there is no clear, abrupt insertion, TPC122 demonstrated progressively stronger and more sustained interaction with the Gram-negative membrane, with an eventually marginally higher contact number compared to the Gram positive membrane (Figure 10A, right-hand side violin plots), reflecting higher membrane affinity and potentially greater perturbation of bilayer organisation in this environment. Per-residue decomposition of each TPC122 residue in contact with all lipids of the NEG (Figure 10B, blue) and POS (orange) indicates slightly higher lipid contacts at specific, N-terminal (Gly1-Phe2) and C-terminal (Phe16-Val22) segments of the peptide, marked with asterisks in the figure, indicating limited and more localised favourable interaction profile for Gram negative relative to Gram positive membranes, suggesting that the preferential antimicrobial activity of TPC122 for *E. coli* relative to *S. aureus* (Table 1, indicated by low MIC) is at least initially driven by close contacts between the two termini of TPC122 with the *E. coli* membrane.

**Figure 10.**
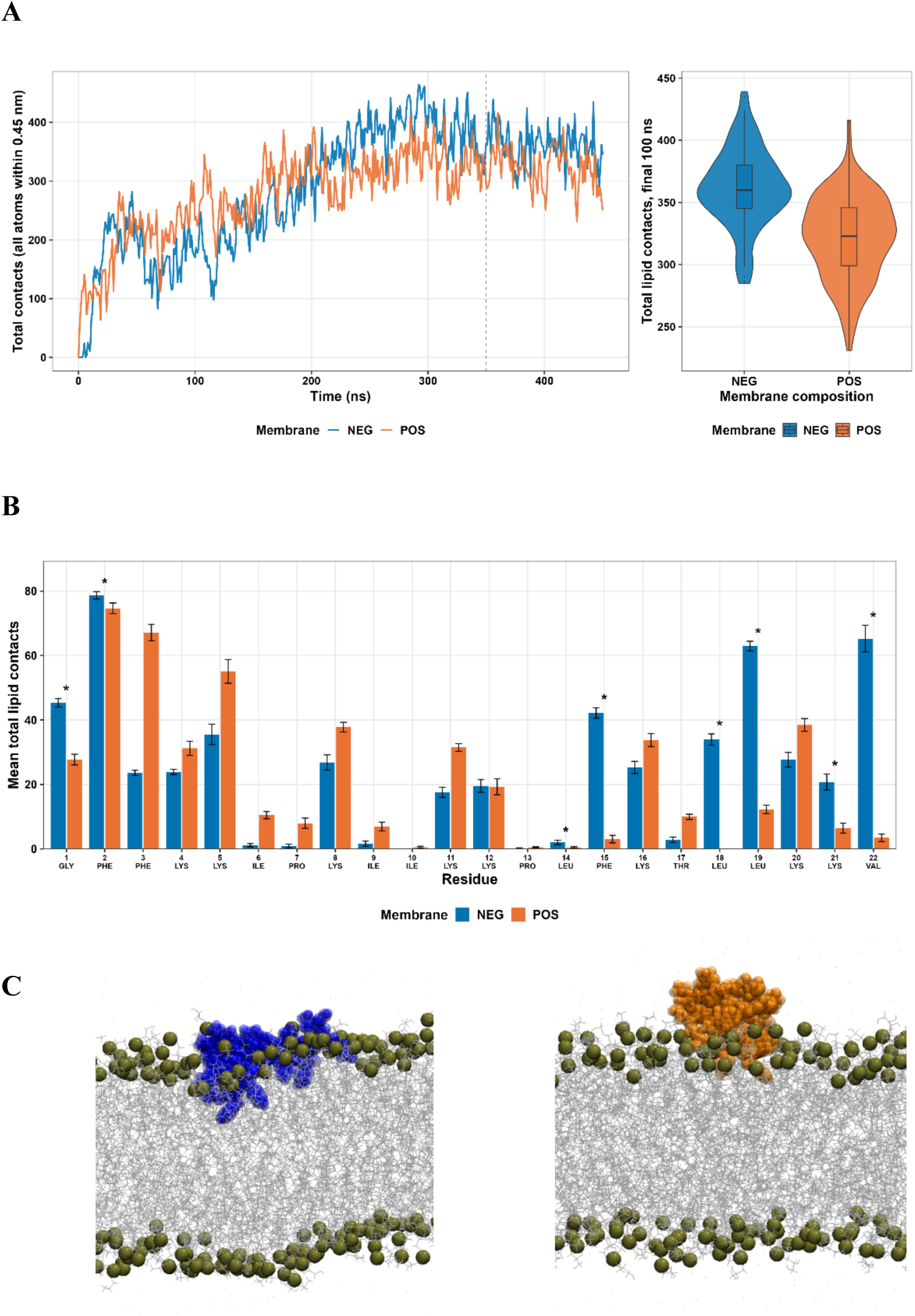
MD simulations of TPC122 at model Gram negative (NEG) and positive (POS) membranes. (A) Time series of interatomic contacts within 0.45 nm between TPC122 peptide and model NEG (blue) and POS (orange) membranes. (B) Per-residue bar charts of peptide-membrane contacts, with averages taken over the final 100 ns. Error bars are five-block-averaged standard errors over the relevant trajectory segment. (C) Graphical snapshots of TPC122 at NEG (blue) and POS (orange) membranes, with bronze spheres representing lipid phosphates, indicating the general extent of membrane insertion.

Secondary structure analysis supports generally higher ellipticity (222 nm) for TPC122 in the presence of a Gram-negative membrane than a Gram-positive membrane for a major portion of the trajectory (Figure 11A; up to ∼300 ns), consistent with increased α-helical content (comparatively illustrated in Figure 11C). This observation was reinforced by the percentage helicity profile, particularly at residues Leu12 to Lys14 (Figure 11B). Taken together, these results indicate that TPC122 preferentially adopts a helical conformation in Gram-negative membranes, accompanied by sustained lipid engagement and insertion for enhanced activity against *E. coli* (Table 1).

**Figure 11.**
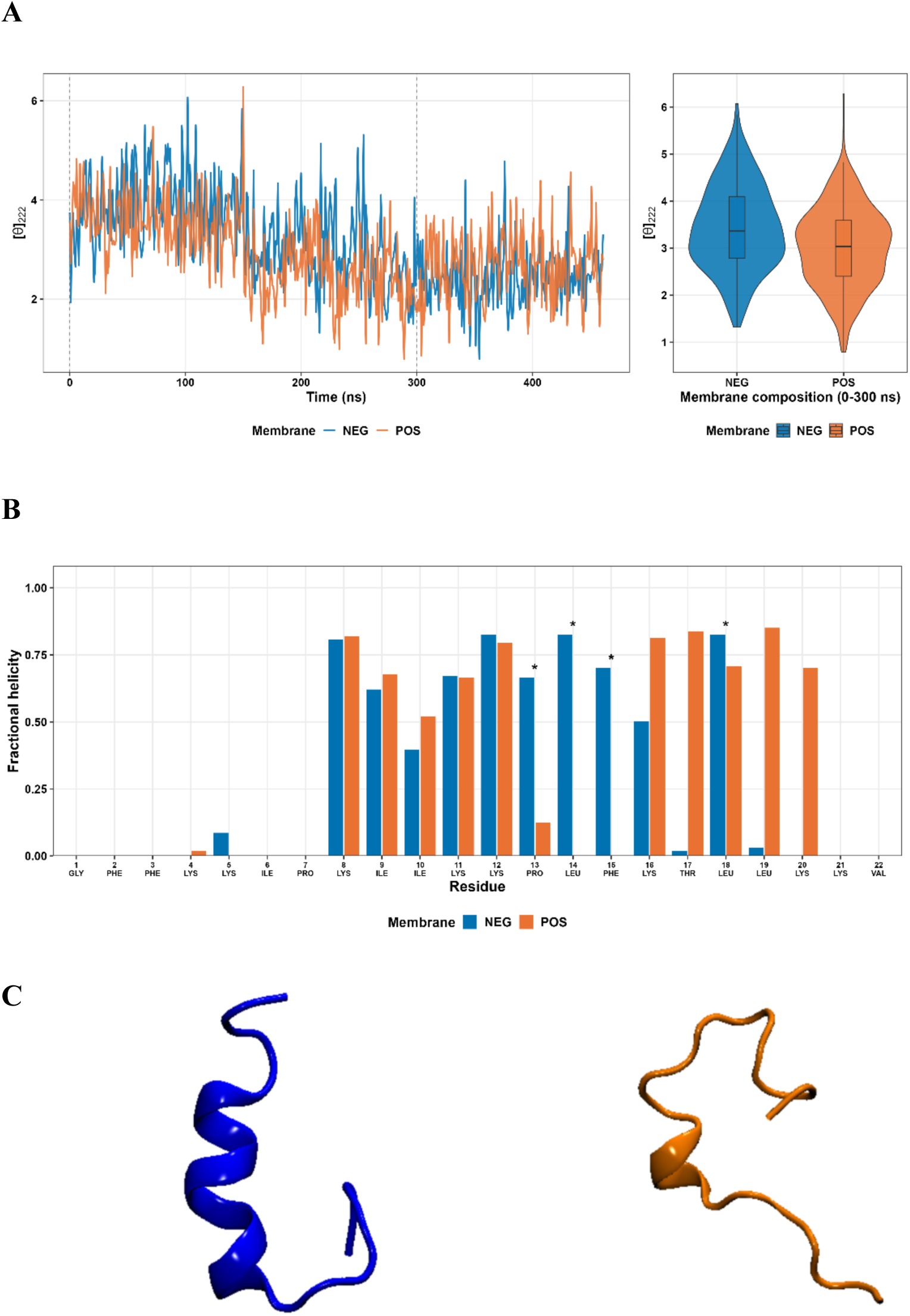
MD simulations of TPC122 at model Gram negative (NEG) and positive (POS) membranes. (A) Time series of ellipticity at 222 nm for TPC122, indicating helicity. (B) Per-residue bar charts of fractional helicity, averaged over the final 100 ns. (C) Graphical snapshots of TPC122 at NEG (blue) and POS (orange) membranes, indicating the general extent of helical content for the peptide at each membrane.

## 4. Conclusions

Based on our recently report on pardaxin (1–22), it possesses greater antibacterial activity and membrane activity against a range of Gram-positive and Gram-negative bacteria, including *Escherichia coli* and *Staphylococcus aureus*. The mechanistic investigation showed pardaxin (1-22) primarily operated through membrane destabilisation and disruption mechanisms. To further identify more pardaxin variants, this work applied computational evolutionary guided deep learning approach to differentiate complex activity-structure relationships over traditional physio-chemical descriptors. Together with our experimental validation, this work also showed that the combination of deep learning methods with specific AMP design considerations to develop novel pardaxin analogues which exhibited selective potency against S. aureus and E. coli, respectively. This highlighted the potential of ESMC as a key technology in the future development of peptide-based therapeutics. The MD simulations and experimental assays further confirmed 10 selected peptides being structurally stable and possessing antimicrobial properties. Overall, our findings highlighted a promising integrated AI-driven workflow which leverages computational prediction, identifying a small subset of highly-likely-potent candidates for subsequent experimental characterisation, in screening a range of novel antimicrobial peptides with enhanced therapeutic value.

## 5. Methods

### 5.1 Peptide descriptor analysis and principal component analysis (PCA)

Peptide descriptor analysis was performed in python [25] using the Peptides package for python, a fork of the peptides package in R [26], to characterize a computationally generated library of sequence variants derived from the parent pardaxin. For each sequence, a comprehensive panel of physicochemical and structural descriptors was calculated, including length, molecular weight, net charge at physiological pH, isoelectric point, hydrophobicity indices, aliphatic index, Boman index, instability index, and amino acid composition–based metrics, as implemented in the Peptides package [26]. Principal component analysis (PCA) was then conducted in python [25] to enable dimensionality reduction and visualization of variance across the descriptor space. Sequences were sampled based on the position of the point in reference to the distribution of the points and their position to known pardaxin activities, combined with distinct selected substitutions which were used to test general AMP theoretical hypothesis, such as increased positive charge, increased hydrophobicity and the alteration of select proline residues at position 7 and 13.

### 5.2 Molecular dynamics simulations

All molecular dynamics systems were constructed and simulated to investigate the adsorption of TPC113 and TPC122 onto model bacterial inner membranes. Initial three-dimensional structures of both peptides were predicted using AlphaFold 3 [27, 28]. Membrane–peptide systems were prepared using the CHARMM-GUI Membrane Builder server [29, 30]. Two symmetric bilayers were generated to approximate bacterial inner membrane compositions: a Gram-negative model consisting of a 7:3 molar ratio of 1-palmitoyl-2-oleoyl-sn-glycero-3-phosphocholine (POPC) and 1-palmitoyl-2-oleoyl-sn-glycero-3-phosphoglycerol (POPG), and a Gram-positive model composed of a 9:1 molar ratio of POPG and 1,1’,2,2’-tetraoleoyl cardiolipin (POCL2) [31, 32]. Each peptide was positioned with its center of mass approximately 10 Å above the membrane phosphate plane and oriented roughly parallel to the bilayer surface to allow spontaneous adsorption during simulation. Systems were solvated using the CHARMM-modified TIP3P water model [33, 34], and counterions were added to ensure overall electroneutrality. All simulations employed the CHARMM36m force field for proteins [35] together with the CHARMM36 lipid force field [36]. MD simulations were performed using GROMACS version 2024.3 [37]. Following steepest descent energy minimization, systems were equilibrated using the standard multi-step CHARMM-GUI protocol with gradually released positional restraints. Production simulations were conducted in the NPT ensemble for 450 ns per system at 310 K and 1 bar, using a Nosé–Hoover thermostat [38, 39] and a semi-isotropic Parrinello–Rahman barostat [40]. Long-range electrostatic interactions were treated with the particle mesh Ewald (PME) method [41], while short-range van der Waals interactions were truncated at 10–12 Å with force switching consistent with CHARMM36. Covalent bonds involving hydrogen atoms were constrained using the LINCS algorithm [42], with a 2 fs integration timestep. Four independent systems were simulated in total, corresponding to TPC113 and TPC122 interacting with each membrane composition, each run for 450 ns, and trajectories were subsequently analyzed using GROMACS analysis tools. Peptide structures were visualised using VMD and ChimeraX [43, 44].

### 5.3 Peptide synthesis and purification

TPC 101, TPC113, TPC 118 and TPC 122 were chemically synthesised using Rink Amide resin by Fmoc/tBu solid-phase methods as previously described [15, 16]. Standard Fmoc-chemistry was used throughout with Rink Amide 4-Methylbenzhydrylamine (MBHA) polystyrene resin (0.1 mmol, loading capacity 0.336 mmol/g, GL. Biochem Ltd., swelling in DMF for 10 min), a coupling condition of a 4-fold molar (0.4 mmol) excess of the Fmoc-protected amino acids in the presence of 4-fold Oxyma Pure (0.4 mmol) and 4-fold DIC (0.4 mmol), and a deprotection condition of 20% piperidine on a peptide synthesizer. After the synthesis, the peptides were cleaved from the solid support resin with trifluoroacetic acid (TFA) in the presence of water and TIPS as the scavenger (ratio 95:2:3) for 2 h at room temperature, followed by filtration to remove the resin. The crude peptide products were obtained with nitrogen evaporation and precipitated in ice-cold diethyl ether and washed three times. The crude peptides were then purified with a C18 column by reversed-phase high performance liquid chromatography (RP-HPLC) in water and acetonitrile containing 0.1% TFA with the gradient 0-100% of buffer B (acetonitrile) at 10 mL/min. The final products were then characterised by both RP-HPLC and Mass Spectrometry as stated in supporting information.

### 5.4 Antimicrobial activity assay

The minimum inhibitory concentrations (MICs) were determined using a broth microdilution method as previously reported [45]. Stock solutions were serially diluted twofold in cation-adjusted Mueller–Hinton broth (CMHB). Equal volumes of each dilution and bacterial suspension were combined in 96-well plates, yielding a final bacterial concentration of 1 × 10^6^ CFU/mL per well. Plates were incubated overnight at 37 °C, and bacterial growth was assessed by measuring optical density at 600 nm (OD600). The MIC was defined as the lowest compound concentration that completely prevented bacterial growth.

### 5.5 Scanning electron microscopy

Scanning electron microscopy (SEM) was employed to investigate peptide-induced morphological changes in bacterial cells. Briefly, overnight bacterial cultures were grown in CMHB medium at 37 °C and adjusted to 1 × 10⁹ CFU/ mL. Equal volumes of the bacterial suspension and peptide solutions were mixed in a 96-well plate to achieve final peptide concentrations of 4, 8, and 16 μg/mL, followed by incubation at 37 °C for 90 min. Subsequently, 20 μL of each sample was spotted onto sterile glass slides and allowed to air-dry. The samples were then fixed with 2.5% glutaraldehyde for 1 h at room temperature, washed with PBS, and dehydrated through an ascending ethanol series (30–100%). After overnight drying, slides were sputter-coated with platinum for 30 s using an SC7640 sputter coater (Quorum Technologies, UK). Imaging was performed on a Hitachi SU7000 SEM at 3 kV, and micrographs were captured at multiple magnifications and analysed using ImageJ.

### 5.6 Outer membrane permeabilisation assay

Test peptides were dissolved in PBS and serially diluted across a 96-well plate according to the desired concentration range. N-Phenyl-1-naphthylamine (NPN) was prepared at 0.5 mM in acetone and diluted with PBS to achieve a final concentration of 10 µM in each well after adding bacterial suspensions and peptide solutions, with bacterial density adjusted to 5 × 10⁹ CFU/mL. Immediately after mixing, fluorescence was measured using a microplate reader at an excitation wavelength of 377 nm and an emission wavelength of 420 nm, with results expressed as relative fluorescence units (RFU). This method enables sensitive evaluation of peptide-induced membrane permeabilisation and provides reliable fluorescence data for subsequent quantitative analysis.

### 5.7 Inner membrane permeabilisation assay

Freshly cultured *E. coli* ATCC 25922 and ATCC 35218 were harvested by centrifugation to remove the growth medium and washed twice with PBS to eliminate residual nutrients. The bacterial pellets were then resuspended in PBS containing ONPG, and 100 µL of this suspension was mixed with 100 µL of serially diluted peptides prepared in the same buffer, resulting in a final bacterial density of 2 × 10⁹ CFU/mL and an ONPG concentration of 1.5 mM in the assay. Absorbance at 420 nm was monitored continuously over 1 hour, with measurements recorded every 5 minutes.

### 5.8 Cytoplasmic membrane permeabilisation assay

Membrane permeabilisation of *S. aureus* was assessed using SYTO Green, a fluorescent dye that selectively enters cells with compromised membranes. Bacterial suspensions at 1 × 10⁹ CFU/mL were incubated with the dye for 10 minutes, after which peptides were added at final concentrations of 8, 16, 32, and 64 µg/mL, with the SYTO Green concentration adjusted to 0.5 µM. Fluorescence intensity, indicative of membrane disruption, was monitored every 2 minutes using a microplate reader with excitation and emission wavelengths of 485 nm and 520 nm, respectively.

## Author contributions

This study was conceived by W.L. and A.H., who conceptualised the research topic and completed the manuscript. Experiments were prepared and performed by T.H., M.P., K.D., A.H. (Computational work), T.Z. A.S. (Biological testing), P.P., A.M., W.L. (Peptide Chemistry). All authors contributed to the project design, manuscript drafting and editing.

## Data availability

The data supporting this article have been included as part of the Supplementary Information.

## Supplementary Information

Supplemental Information, including characterisation of candidate peptides, PCA analysis, and peptide-induced membrane permeabilisation assays, can be found with this article.

## Acknowledgement

We would like to acknowledge La Trobe University Bioimaging and Comprehensive Proteomics Platform, Proteomics and Metabolomics Platform for the provision of instrumentation, training and technical support. This work was funded in part by a National Health and Medical Research Council Emerging Investigator Grant to WL (APP2018256) and Ramaciotti Foundations (WL). We gratefully acknowledge the computational resources provided by the National Computational Infrastructure (NCI), which is supported by the Australian Government’s National Collaborative Research Infrastructure Strategy (NCRIS) We also acknowledge the Pawsey Supercomputing Research Centre, a world-class high-performance computing facility in Perth, Western Australia, which is funded by the Australian Government and the Government of Western Australia under NCRIS. Additionally, we thank the RMIT AWS Cloud Supercomputing Hub (RACE Hub) for providing access to scalable cloud computing resources through their collaboration with Amazon Web Services. M.P. is supported by a ACAMI PhD scholarship. We thank Yujie Zhu’s assistance during peptide preparation.

