## Supplemental document for "Computational evolutionary approach to generate antimicrobial peptides"

**Table of Contents**


1. *Characterisation of candidate peptides* ........................................ *S2*
2. *PCA analysis* ........................................ *S5*
3. *Peptide-induced outer membrane permeabilisation in E. coli* ........................................ *S10*
4. *Peptide-induced inner membrane permeabilisation in E. coli* ........................................ *S11*
5. *Peptide-induced cytoplasmic membrane permeabilisation in S. aureus* ........................................ *S12*

### 1. Characterisation of candidate peptides

*Table S1. The sequence information of synthesised peptides used in this report*

| Peptide | Sequences | Calculated MW | Observed MW |
|---|---|---|---|
| TPC101 (Pardaxin)[a] | GFFALIPKIISSPLFKTLLSAV-$NH_2$ | 2465.64 | 1201.4 $[M+2H]^{2+}$ |
| TPC109 | GFFALIPKIISSHLFKTLLSAV-$NH_2$ | 2401.93 | 1218.7 $[M+2H]^{2+}$ |
| TPC110 | GFFALIPKIISSFLFKTLLSAV-$NH_2$ | 2411.96 | 1206.8 $[M+2H]^{2+}$ |
| TPC111 | GFFALIWKIISSPLFKTLLSAV-$NH_2$ | 2451 | 1226.35 $[M+2H]^{2+}$ |
| TPC112 | GFFALIPKIISSPLFKTLLHAV-$NH_2$ | 2411.97 | 1206.85 $[M+2H]^{2+}$ |
| TPC113 | GFFAWIPKIISSPLFKTLLSAV-$NH_2$ | 2434.6 | 1218 $[M+2H]^{2+}$ |
| TPC114 | GFFFLIPKIISSPLFKTLLSAV-$NH_2$ | 2438 | 1218.1 $[M+2H]^{2+}$ |
| TPC115 | GFFALIVKIISSPLFKTLLSAV-$NH_2$ | 2363.92 | 1182.5 $[M+2H]^{2+}$ |
| TPC117 | GFFALIPKIISRPLFKTLLSAV-$NH_2$ | 2431.05 | 1216.06 $[M+2H]^{2+}$ |
| TPC118 | GFFALIPKKISSPLFKTLLSAV-$NH_2$ | 2376.69 | 1189.06 $[M+2H]^{2+}$ |
| TPC119 | GFFALIPKIISSRLFKTLLSAV-$NH_2$ | 2420.989 | 1211.24 $[M+2H]^{2+}$ |
| TPC 120 | GFFALIPKIISSKLFKTLLSAV-$NH_2$ | 2392.976 | 1197.18 $[M+2H]^{2+}$ |
| TPC121 | GFFRLIPKIISSPLFKTLLSAV-$NH_2$ | 2447.027 | 1224.28 $[M+2H]^{2+}$ |
| TPC122 | GFFKKIPKIIKKPLFKTLLKKV-$NH_2$ | 2614.411 | 1308.12 $[M+2H]^{2+}$ |
| TPC123 | GFKKLIPKIIKKPLFKTLLKKV-$NH_2$ | 2580.394 | 860.9 $[M+3H]^{3+}$ |

a: The peptide TPC101 has been reported previously[1, 2].

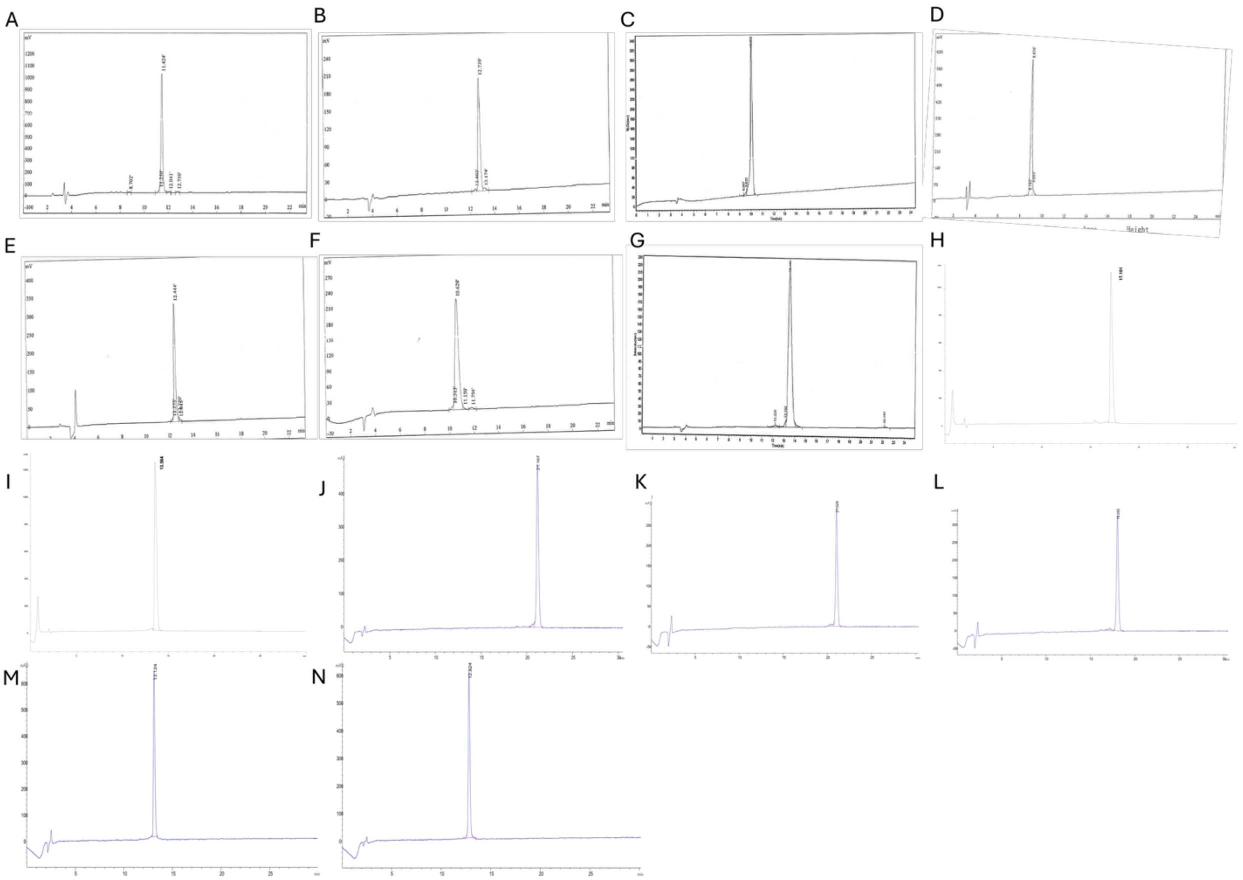


***Figure S1.*** *Analytical HPLC chromatograms of synthetic peptides. HPLC conditions: C18 column (5 µm, 4.6 × 250 mm), linear gradient of 0–100% acetonitrile (containing 0.1% TFA) in water (containing 0.1% TFA) over 30 min. (A) TPC109; (B) TPC110; (C) TPC111; (D) TPC112; (E) TPC113; (F) TPC114; (G) TPC115; (H) TPC117; (I) TPC118; (J) TPC119; (K) TPC120; (L) TPC121; (M) TPC122; (N) TPC123.*

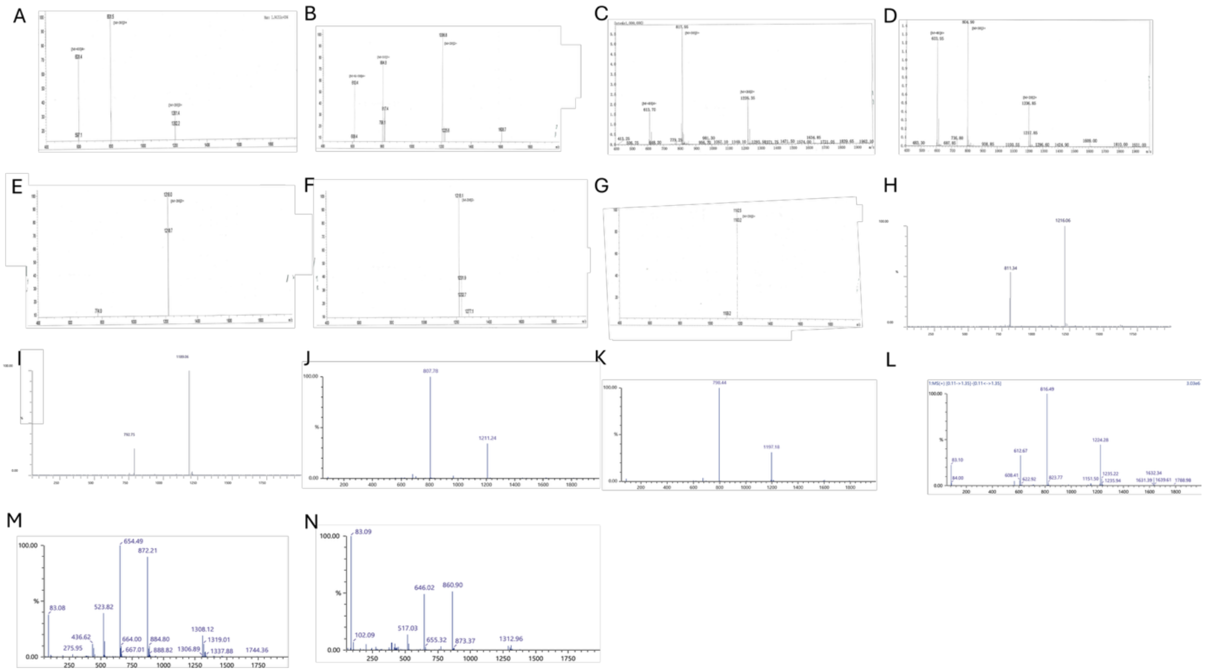


*Figure S2. Mass spectrometry characterization of synthetic peptides. (A) TPC109, calculated MS: 2401.93Da, found MS:* m/z 1226.35 $[M+2H]^{2+}$*; (B) TPC110, calculated MS: 2411.96 Da, found MS:* m/z 1206.8 $[M+2H]^{2+}$*; (C) TPC111, calculated MS: 2451 Da, found MS:* m/z 1226.35 $[M+2H]^{2+}$*; (D) TPC112, calculated MS: 2411.97 Da, found MS:* m/z 1206.85 $[M+2H]^{2+}$*; (E) TPC113, calculated MS: 2434.6 Da, found MS:* m/z 1218 $[M+2H]^{2+}$*; (F) TPC114, calculated MS: 2438Da, found MS:* m/z 1218.1 $[M+2H]^{2+}$*; (G) TPC115, calculated MS: 2363.92 Da, found MS:* m/z 1182.5 $[M+2H]^{2+}$*; (H) TPC117, calculated MS: 2431.05 Da, found MS:* m/z 1216.06 $[M+2H]^{2+}$*; (I) TPC118, calculated MS:* 2376.69 *Da, found MS:* m/z 1189.06 $[M+2H]^{2+}$*;; (J) TPC119, calculated MS: 2420.989 Da, found MS:* m/z 1211.24 $[M+2H]^{2+}$*; (K) TPC120, calculated MS: 2392.976 Da, found MS:* m/z 1197.18 $[M+2H]^{2+}$*; (L) TPC121, calculated MS: 2447.027 Da, found MS:* m/z 1224.28 $[M+2H]^{2+}$*; (M) TPC122, calculated MS: 2614.411Da, found MS:* m/z 1308.12 $[M+2H]^{2+}$*; (N) TPC123, calculated MS: 2580.394 Da, found MS:* m/z 860.9, $[M+3H]^{3+}$. *The observed masses were in good agreement with the calculated values, confirming the identity of all synthesized peptides.*

### 2. PCA analysis

#### 2.1 PCA analysis for Magainin-2 Variants

Sequence: GIGKFLHSAKKFGKAWVGEIMNS

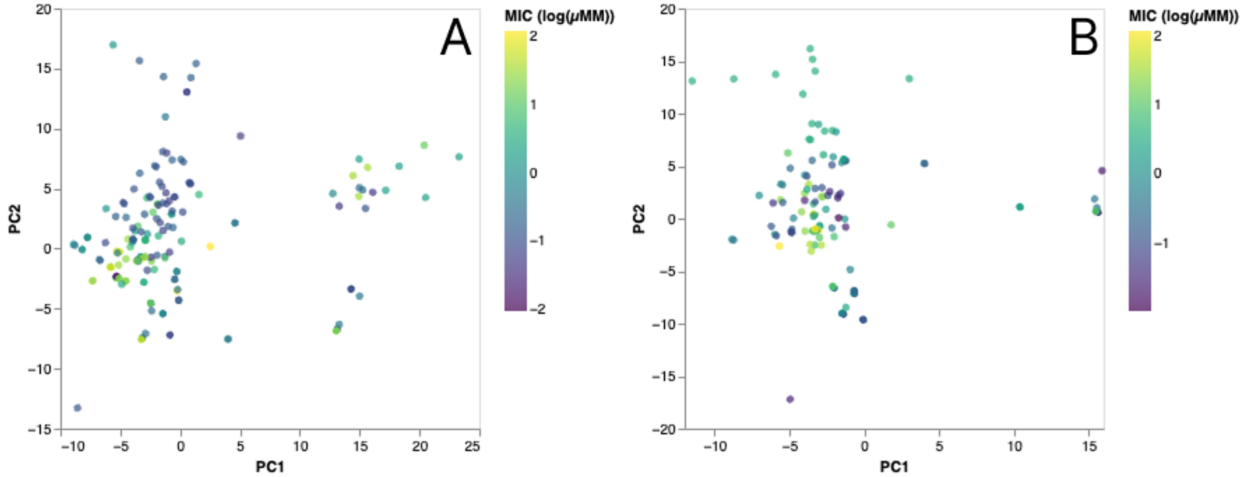


*Figure S3. Scatter plot of Magainin-2 sequences with sequence identity of > 0.8. PCA1 and PC2 are plotted from 112 calculated physiochemical descriptors for variants with known EC activity (A) and known SA activity (B).*

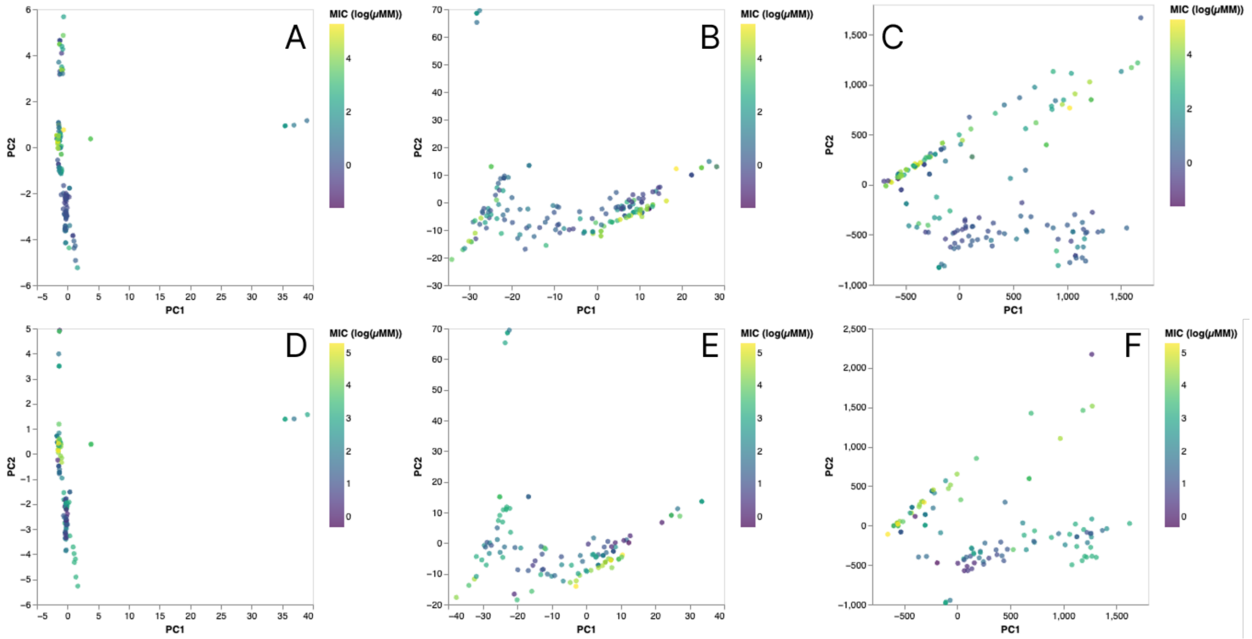


*Figure S4. Scatter plot of Magainin-2 sequences with sequence identity of > 0.8. PCA1 and PC2 are plotted from ESMC embedding descriptors for variants with known EC activity (top) and*

*known SA activity (bottom). (A, D) are the embeddings from layer 0, (B, E) are the embeddings from layer 15 and (C, F) are the embeddings from layer 29.*

### 2.2 PCA analysis for Peptide-P Variants

Sequence: KWKSFLKTFKSAKKTVKHTLLKAISS

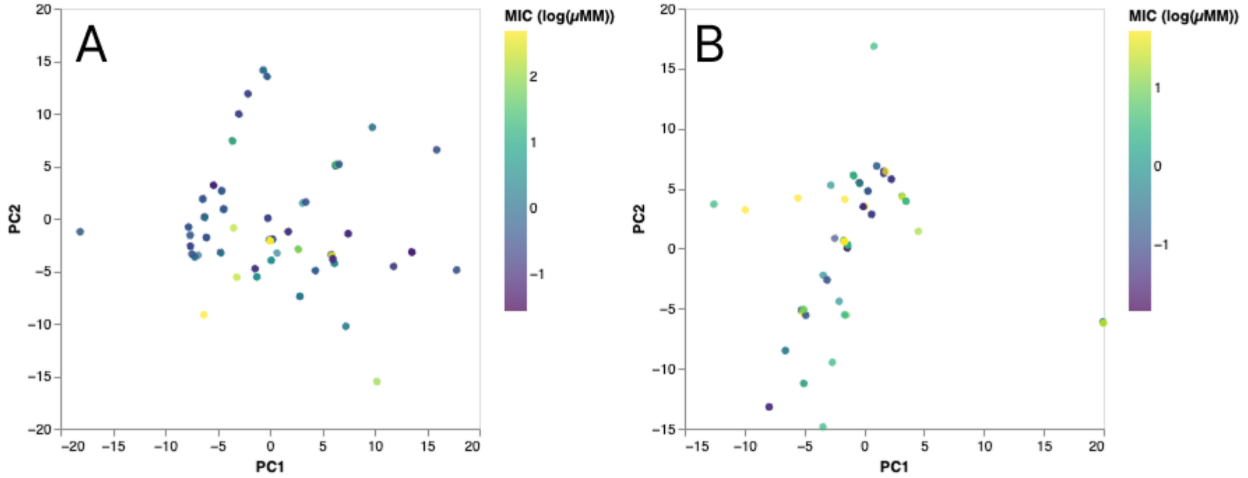


*Figure S5. Scatter plot of Peptide-P sequences with sequence identity of > 0.8. PCA1 and PC2 are plotted from 112 calculated physiochemical descriptors for variants with known EC activity (A) and known SA activity (B).*

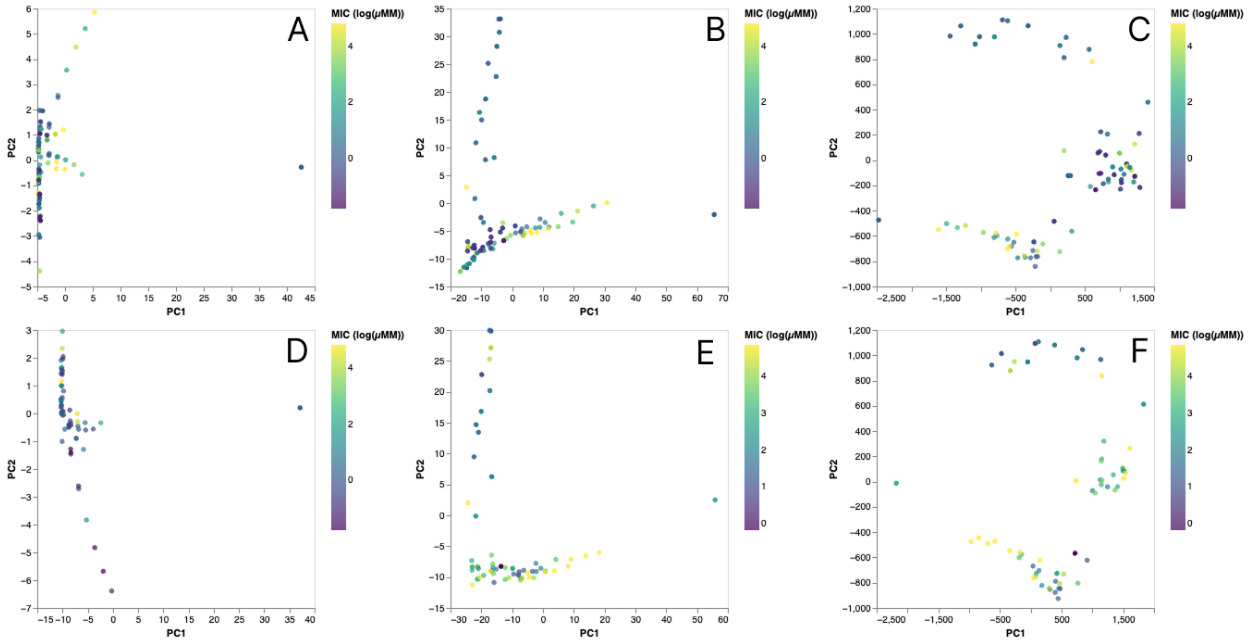

*Figure S6. Scatter plot of Peptide-P sequences with sequence identity of > 0.8. PCA1 and PC2 are plotted from ESMC embedding descriptors for variants with known EC activity (top) and known SA activity (bottom). (A, D) are the embeddings from layer 0, (B, E) are the embeddings from layer 15 and (C, F) are the embeddings from layer 29.*

**2.3 PCA analysis for Oncocin and Pyrrhocoricin Variants**

Sequence: VDKPPYLPRPHPPRRIYNNR

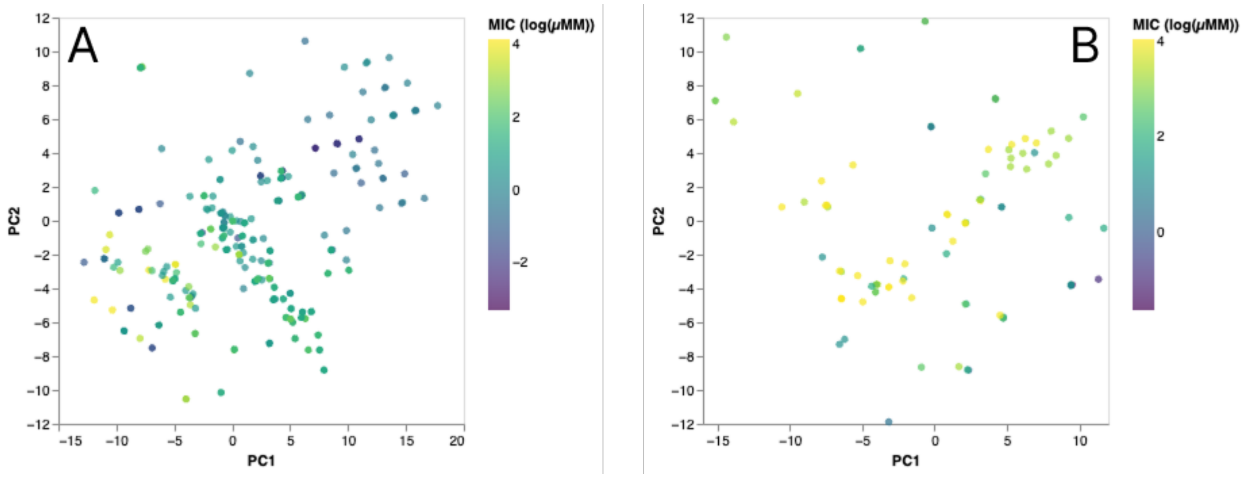


*Figure S7. Scatter plot of oncocin and pyrrhocoricin sequences with sequence identity of > 0.8. PCA1 and PC2 are plotted from 112 calculated physiochemical descriptors for variants with known EC activity (A) and known SA activity (B).*

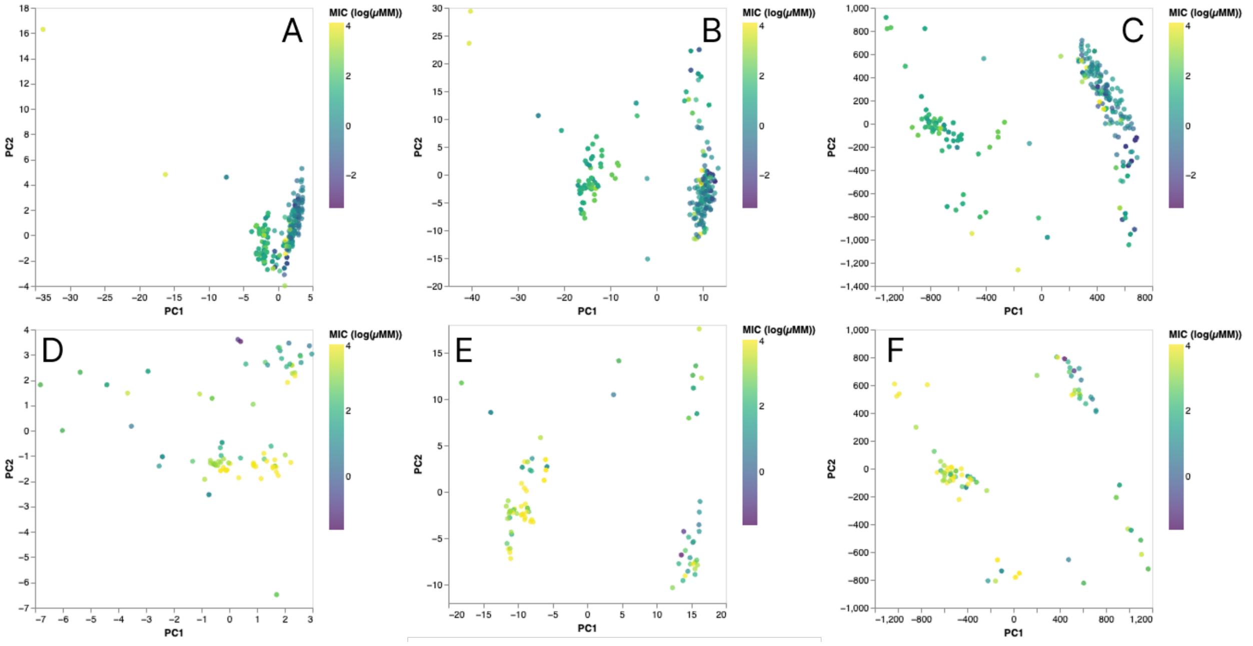


*Figure S8. Scatter plot of oncocin and pyrrhocoricin sequences with sequence identity of > 0.8. PCA1 and PC2 are plotted from ESMC embedding descriptors for variants with known EC activity (top) and known SA activity (bottom). (A, D) are the embeddings from layer 0, (B, E) are the embeddings from layer 15 and (C, F) are the embeddings from layer 29.*

**2.4 PCA analysis for Pardaxin Variants**

Sequence: VDKPPYLPRPHPPRRIYNNR

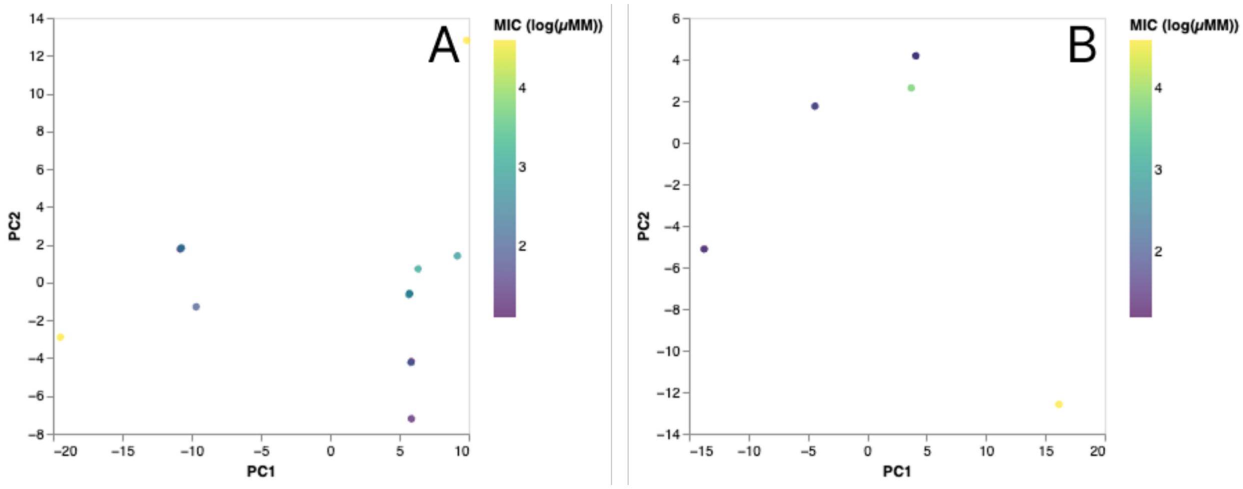

*Figure S9. Scatter plot of pardaxin sequences with sequence identity of > 0.8. PCA1 and PC2 are plotted from 112 calculated physiochemical descriptors for variants with known EC activity (A) and known SA activity (B).*

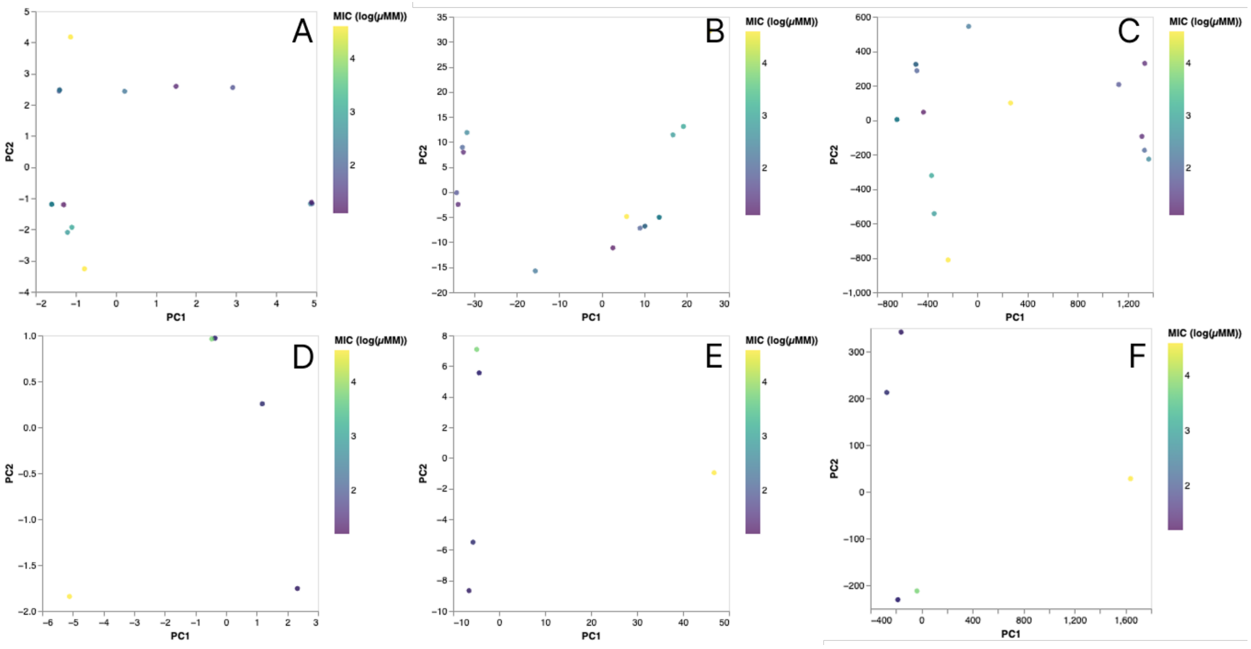


*Figure S10. Scatter plot of pardaxin sequences with sequence identity of > 0.8. PCA1 and PC2 are plotted from ESMC embedding descriptors for variants with known EC activity (top) and known SA activity (bottom). (A, D) are the embeddings from layer 0, (B, E) are the embeddings from layer 15 and (C, F) are the embeddings from layer 29.*

### *3.* Peptide-induced outer membrane permeabilisation in *E. coli*

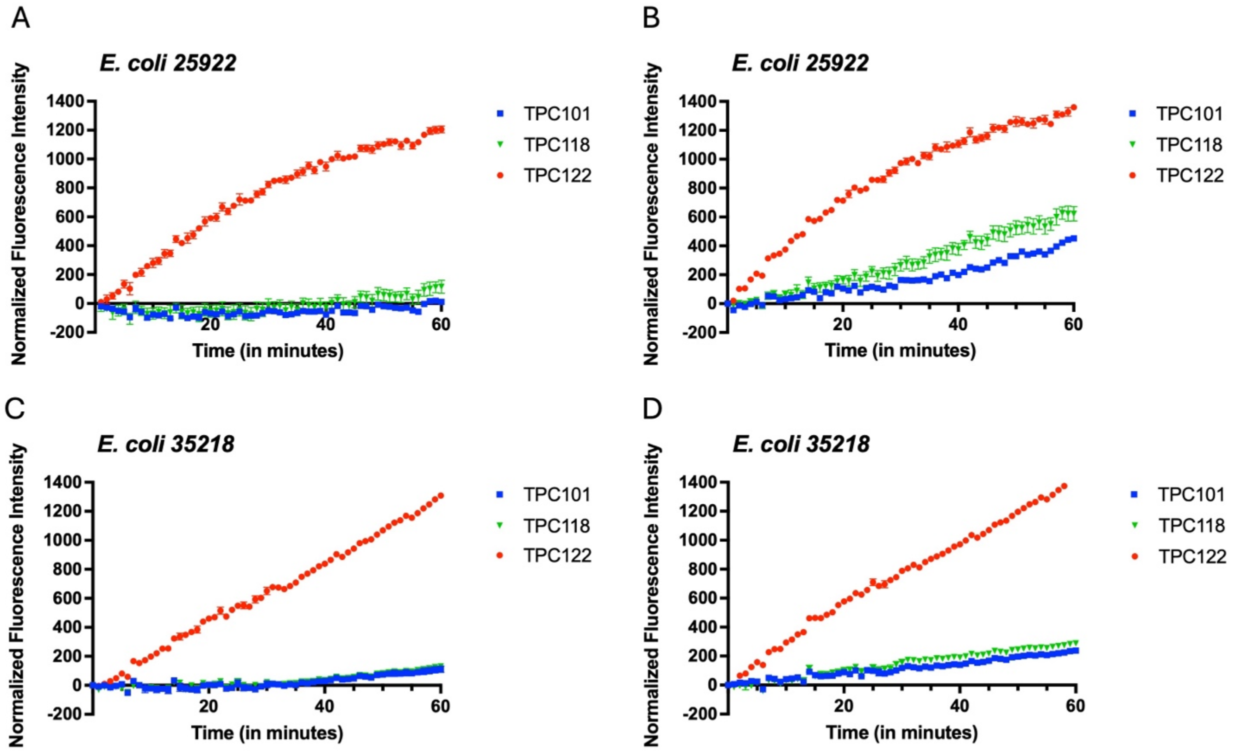


*Figure S11. Peptide-induced outer membrane permeabilisation. NPN fluorescence kinetics were monitored to assess outer membrane permeabilisation following treatment with the indicated peptides. (A–B) E. coli 25922 and (C–D) E. coli 35218 were exposed to 4 and 16 µg/mL, and fluorescence intensity was recorded over time. Enhanced NPN fluorescence indicates increased disruption of the bacterial outer membrane. Data represent mean ± SEM from three independent experiments.*

### *4.* Peptide-induced inner membrane permeabilisation in *E. coli*

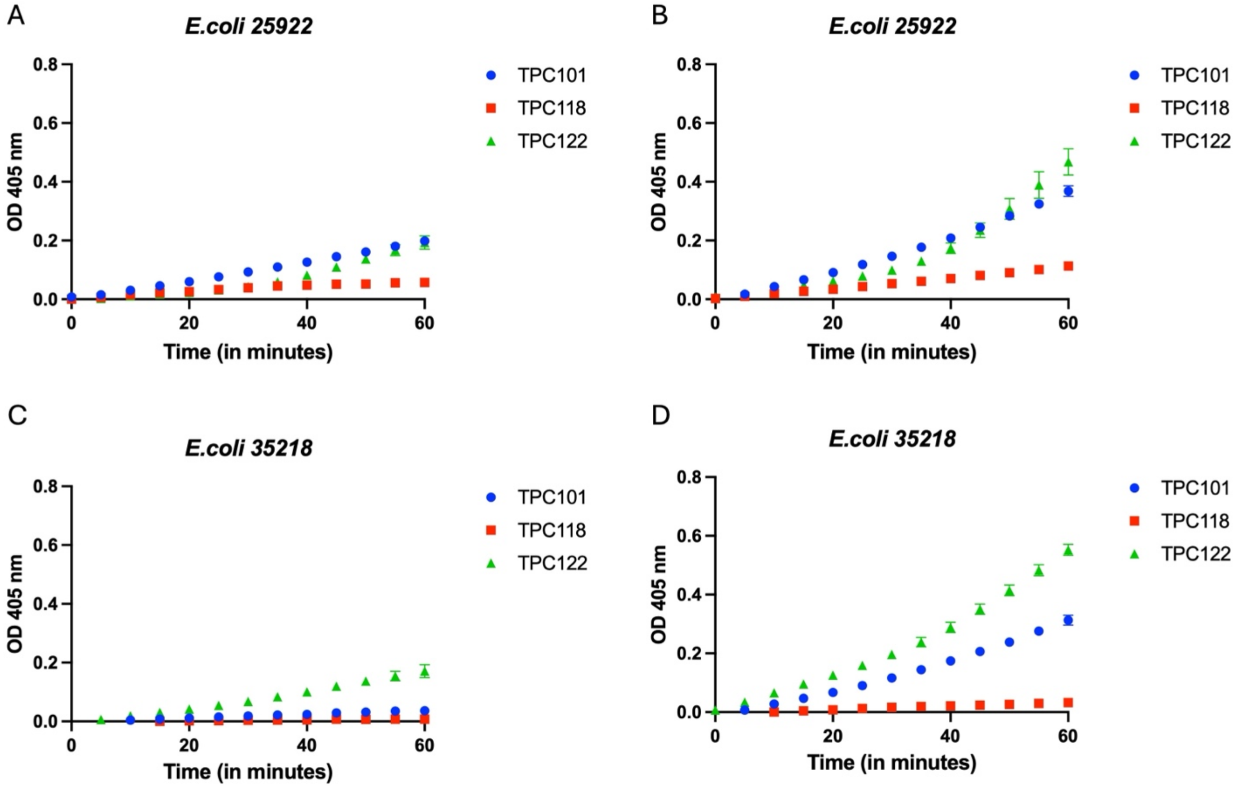


*Figure S12. Peptide-induced inner membrane permeabilisation. ONPG hydrolysis was used to assess cytoplasmic membrane permeabilisation following treatment with the indicated peptides. (A–B) E. coli 25922 and (C–D) E. coli 35218 were exposed to 4 and 16 µg/mL, and the formation of o-nitrophenol was monitored spectrophotometrically over time. Increased absorbance reflects enhanced cytoplasmic membrane permeabilisation, allowing ONPG access to intracellular β-galactosidase. Data represent mean ± SEM from three independent experiments.*

### 5. Peptide-induced cytoplasmic membrane permeabilisation in *S. aureus*

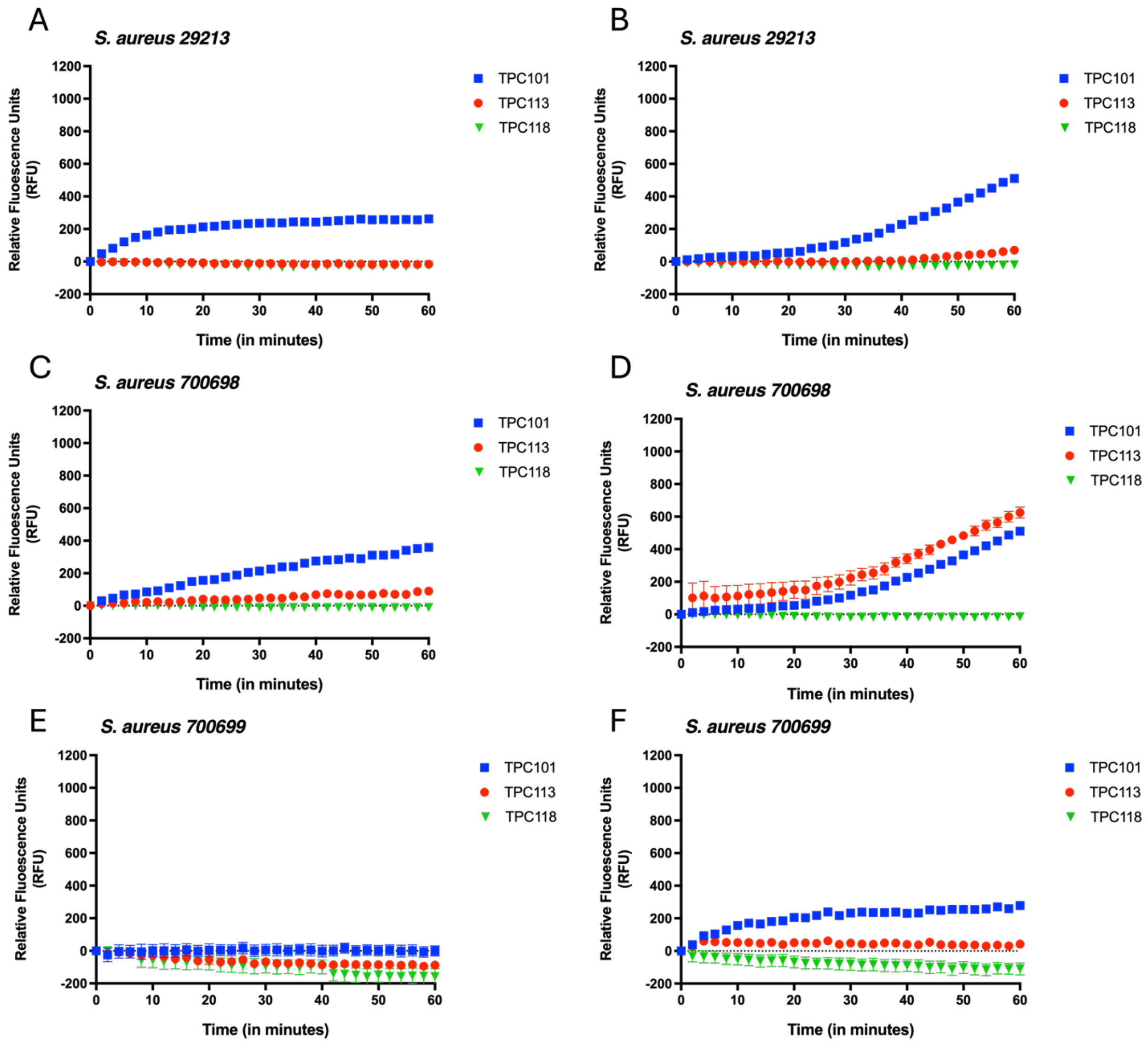


*Figure S12. Peptide-induced membrane permeabilisation in S. aureus. Membrane integrity was assessed by monitoring fluorescence kinetics following treatment with the indicated peptides. (A–B) S. aureus 29213, (C–D) S. aureus 700698, and (E–F) S. aureus 700699 were treated at 4 and 16 µg/mL, and fluorescence intensity was recorded over time. Increased fluorescence reflects enhanced cytoplasmic membrane permeabilisation. Data represent the mean ± SEM from three independent experiments.*
